# Genome-wide methylation data improves dissection of the effect of smoking on body mass index

**DOI:** 10.1101/2020.10.08.329672

**Authors:** Carmen Amador, Yanni Zeng, Michael Barber, Rosie Walker, Archie Campbell, Andrew M. McIntosh, Kathryn L. Evans, David Porteous, Caroline Hayward, James F. Wilson, Pau Navarro, Chris S. Haley

## Abstract

Variation in obesity-related traits has a genetic basis with heritabilities between 40 and 70%. While the global obesity pandemic is usually associated with environmental changes related to lifestyle and socioeconomic changes, most genetic studies do not include all relevant environmental covariates, so genetic contribution to variation in obesity-related traits cannot be accurately assessed. Some studies have described interactions between a few individual genes linked to obesity and environmental variables but there is no agreement on their total contribution to differences between individuals. Here we compared self-reported smoking data and a methylation-based proxy to explore the effect of smoking and genome-by-smoking interactions on obesity related traits from a genome-wide perspective to estimate the amount of variance they explain. Our results indicate that exploiting omic measures can improve models for complex traits such as obesity and can be used as a substitute for, or jointly with, environmental records to better understand causes of disease.

## Introduction

Variation in obesity-related traits such as body mass index (BMI) has a complex basis with heritabilities ranging from 40 to 70%, with the genetic variants detected to date explaining up to 5% of BMI variation^1^. In addition to genetics, studies suggest that the increase in obesity prevalence in recent decades is linked to environmental causes, such as dietary changes and a more sedentary lifestyle^2,3,4,5^. The fact that all relevant environmental effects have not been accounted for in genetic studies has potentially reduced GWAS power to detect susceptibility variants. On top of this, several studies suggest that gene-by-environment interactions also play an important role in obesity and other complex traits ^2,6,7,8,9,10^ and many researchers are focusing on finding interactions between specific genes and certain environments. Genotype-by-age interactions and genotype-by-sex interactions have also been detected for several health-related traits^10,11,12^. Recently, when performing GWAS on traits like BMI, lipids, and blood pressure, several studies have stratified their samples on the basis of smoking status or have explicitly modelled interactions leading to identification of new genetic variants associated with those traits ^13,14,15^. Some studies have attempted to quantify the overall contribution of genetic interactions with smoking. Robinson, et al.^12^ estimated them to explain around 4% of BMI variation in a subset of unrelated UK Biobank samples. In contrast, also in UK Biobank, using a new approach that only requires summary statistics, Shin & Lee^17^ estimated the contributions of the interactions to be much smaller: 0.6% of BMI variation.

In this study, we aim to estimate the contribution of smoking and its interaction with genetic variation to obesity variation, using self-reported measures of smoking and a methylomic proxy of smoking exposure. We hypothesised that use of a proxy, rather than self-reported smoking, and fitting gene by smoking interactions would lead to more a more accurate model. DNA methylation is an epigenetic mark that can be affected by genetics and environmental exposures^18,19,20,21,22,23^. Variation in methylation is correlated with gene expression, plays a crucial role in development, in maintaining genomic stability^24,25,26^, and has been associated with disease^27,28,29,30,31^ and aging^32,33^. Epigenome-wide association analyses (EWAS) have identified multiple associations between DNA methylation levels at specific genomic locations and smoking^19,34,35,36^. These so-called *signatures* of smoking in the epigenome can help discriminate the smoking status of the individuals in a cohort^20^, and, if sufficiently accurate, could be an improvement on self-reported measures, by adding information not captured (accurately) in the self-reported measure, such as passive smoking or real quantity of tobacco smoked.

Here, we aim to estimate the contribution to obesity variation of smoking and its interaction with genetic variation in two different cohorts, using self-reported measures of smoking and a methylomic proxy for smoking. Thus, we measured the contribution of smoking-associated methylation signatures and genome-by-methylation interactions to trait variation. We performed analyses in both sexes jointly and independently and also including genome-by-smoking-by-sex interactions, and we showed that omics data can be exploited as proxies for environmental exposures to improve our understanding of complex trait architecture. We observed that using an appropriate set of CpG sites, methylation can be used to model trait variation associated with smoking, and genome-by-smoking interactions suggesting potential applications for better prediction and prognosis of complex disease and expanding these modelling approaches to other environments and traits.

## Results

The aim of this work was to explore the influence of smoking and genome-by-smoking interactions on trait variation, modelling them from self-reported information and using DNA methylation in both sexes jointly and separately. We used a variance component approach to fit a linear mixed model including a set of covariance matrices representing: two genetic effects (G: common SNP-associated genetic effects and K: pedigree-associated genetic effects not captured by the genotyped markers at a population level; the inclusion of matrix K in the analyses allows to use the related individuals in the sample), environmental effects reflecting impact of smoking (modelled as fixed or random effects), and genome-by-smoking effects (GxSmk) representing sharing of both genetics (G) and environment (smoking, Smk), and we estimated the proportion of variation that each component explained for seven obesity-related measures: weight, body mass index (BMI), waist circumference (waist), hip circumference (hips), waist-to-hip ratio (WHR), fat percentage (fat%), and HDL cholesterol (HDL) as well as height, to serve as a negative control. We defined the environment using either self-reported questionnaire data or its associated methylation signature as a proxy. A summary of the experimental design used in this study is shown in Figure 1. For more detailed information, see Methods.

**Figure 1.**
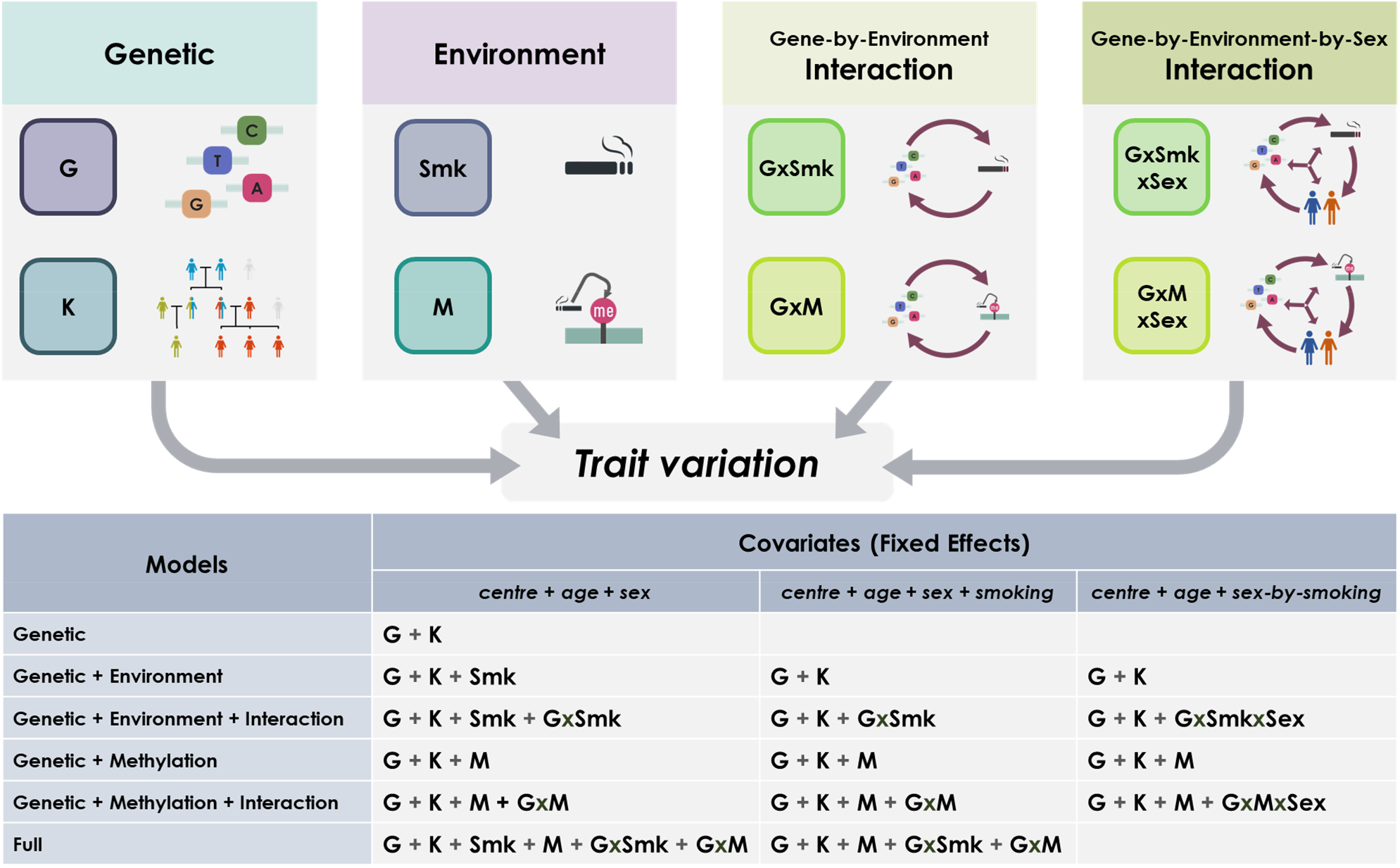
Summary of the experimental design of the study. The panels (above) represent the genetic and environmental components contributing to trait variation and used in the models (table below). Each cell shows the included random effects in each combination of model (row) and fixed effects (columns). G: Genomic, K: Kinship, GxSmk: Genome-by-Smoking, M: Methylation, GxM: Genome-by-Methylation, GxSmkxSex: Genome-by-Smoking-by-Sex, GxMxSex: Genome-by-Methylation-by-Sex. Models applied to different data sets varied depending on data availability.

### Self-reported smoking status

#### Generation Scotland

Figure 2 shows the estimates of the proportion of BMI, fat percentage, and HDL variance explained by different sources included in the linear mixed models in ~18K individuals in Generation Scotland (GS18K). Results for other traits are displayed in Table 1, Supplementary Figure 1, and full details of the analyses for all traits including estimates, standard errors, and log-likelihood ratio tests (LRT) are shown in Supplementary Table 1.

**Figure 2.**
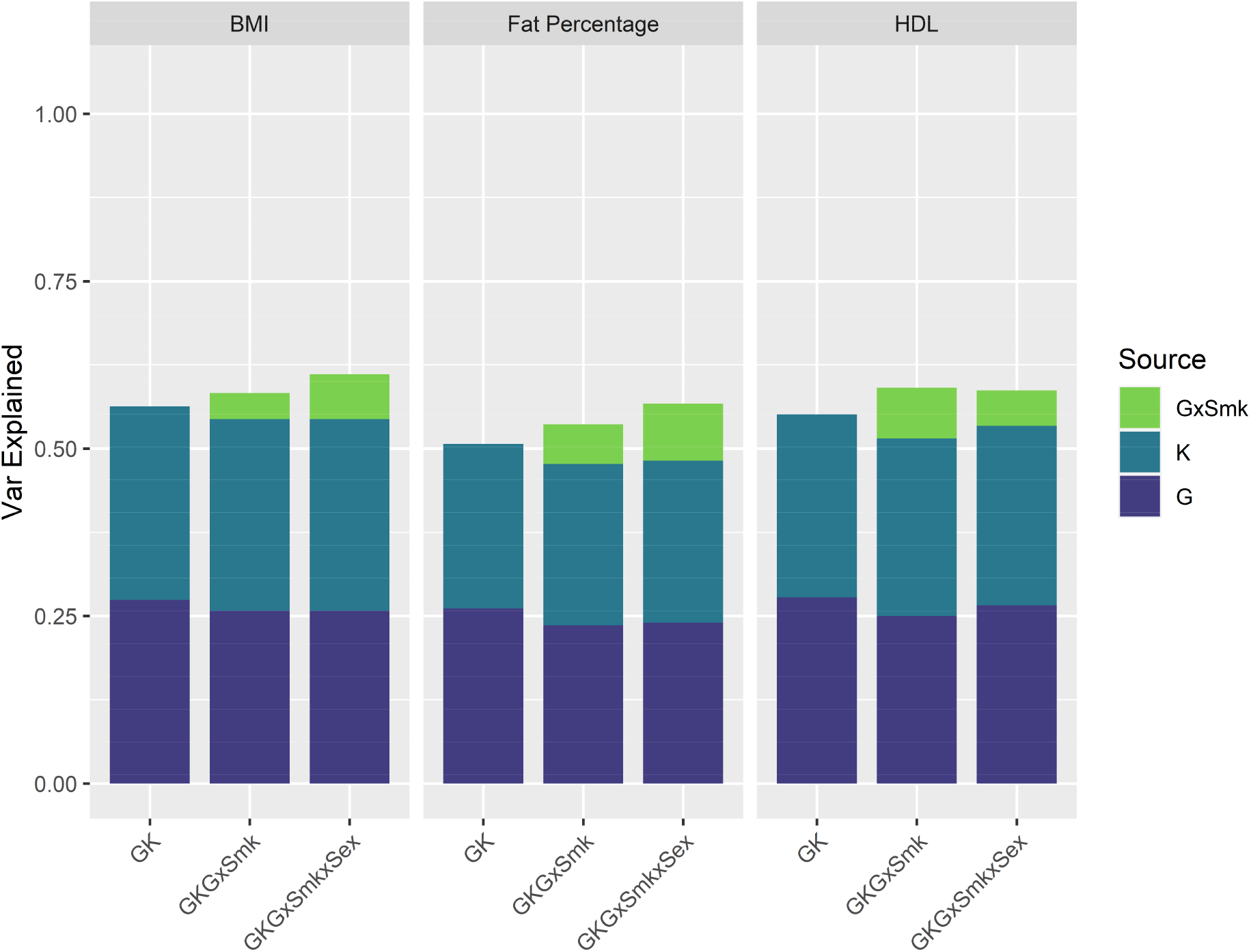
Proportion of trait variation explained by genetic and interaction sources in GS18K. Proportion of BMI, fat percentage, and HDL variance (y-axis) explained by each of the genetic and interaction sources in the corresponding models (x-axis). G: Genomic, K: Kinship, GxSmk: Genome-by-Smoking.

**Table 1.** Summary of interaction results for all cohorts. Results of GKGxSmk model for all traits in GS18K and meta-analysis of the recruitment centre-based sub-cohorts in UK Biobank. The table shows, for each trait, the proportion of the phenotypic variance explained (Var), its standard error (SE), the log-likelihood ratio test P value (LRT P, only for the interaction), the meta-analysis P value (P), for each of the components in the model: Genetic (G), Kinship (K) and genome-by-smoking interaction (GxSmk). Highlighted P values indicate nominally significant results for the GxSmk component.

| Trait | Source | GS18K |  |  | UKB Meta Analysis |  |  |
| --- | --- | --- | --- | --- | --- | --- | --- |
|  |  | Var | SE | LRT P | Var | SE | P |
| Height | G | 0.483 | 0.022 |  | 0.629 | 0.009 |  |
| Height | K | 0.429 | 0.024 |  | 0.328 | 0.006 |  |
| Height | GxSmk | 0.012 | 0.014 | 0.2041 | 0.001 | 0.003 | 0.7640 |
| Weight | G | 0.270 | 0.024 |  | 0.355 | 0.007 |  |
| Weight | K | 0.302 | 0.027 |  | 0.242 | 0.018 |  |
| Weight | GxSmk | 0.049 | 0.021 | 0.0098 | 0.022 | 0.008 | 0.0050 |
| BMI | G | 0.258 | 0.024 |  | 0.318 | 0.008 |  |
| BMI | K | 0.286 | 0.028 |  | 0.236 | 0.021 |  |
| BMI | GxSmk | 0.039 | 0.021 | 0.0336 | 0.025 | 0.007 | 0.0009 |
| Waist | G | 0.181 | 0.024 |  | 0.261 | 0.004 |  |
| Waist | K | 0.313 | 0.028 |  | 0.214 | 0.021 |  |
| Waist | GxSmk | 0.023 | 0.022 | 0.1534 | 0.017 | 0.007 | 0.0119 |
| Hips | G | 0.212 | 0.024 |  | 0.296 | 0.009 |  |
| Hips | K | 0.271 | 0.028 |  | 0.179 | 0.028 |  |
| Hips | GxSmk | 0.027 | 0.023 | 0.1185 | 0.020 | 0.007 | 0.0048 |
| WHR | G | 0.130 | 0.023 |  | 0.217 | 0.005 |  |
| WHR | K | 0.198 | 0.027 |  | 0.151 | 0.013 |  |
| WHR | GxSmk | 0.019 | 0.023 | 0.2011 | 0.012 | 0.006 | 0.0437 |
| Fat% | G | 0.236 | 0.025 |  | 0.301 | 0.006 |  |
| Fat% | K | 0.241 | 0.028 |  | 0.224 | 0.013 |  |
| Fat% | GxSmk | 0.059 | 0.023 | 0.0036 | 0.021 | 0.005 | 0.0000 |
| HDL | G | 0.250 | 0.024 |  |  | NA |  |
| HDL | K | 0.265 | 0.027 |  |  | NA |  |
| HDL | GxSmk | 0.076 | 0.022 | 0.0002 |  | NA |  |

The heritability estimates of all analysed traits (i.e., proportion of the variance captured by G and K matrices together) are consistent with previous estimates in the same cohort^37^. The estimated contributions of smoking status (and the other covariates) to trait variation ranged between 0.35% (for height, assessed as a negative control, as we do not expect to find the same type of effects as with obesity-related measures) and 1.2% (for HDL cholesterol) and are shown in Supplementary Table 2. When included as random effect, smoking explained between 0.1% (for height) and 2.5% (for HDL cholesterol) of trait variation (Supplementary Table 1). Our models identified significant genome-by-smoking interactions for weight, BMI, fat percentage and HDL cholesterol (with log-likelihood ratio tests showing that the models including the interaction were significantly better), explaining between 4 and 8% of trait variation (Table 1), similar to the values of Robinson et al.^12^ for BMI. When the interactions included sex (genome-by-smoking-by-sex interactions) the component was significant for all traits, and explained variance ranging between 2-9% (Supplementary Table 2).

#### UK Biobank

We sought to replicate the results observed in Generation Scotland with data from the UK Biobank cohort (UKB). Analyses were run in four sub-cohorts for computational reasons (G1, G2, G3 and G4, grouping individuals in geographically close recruitment centres; for more information see Methods and Supplementary Table 3), with the two sexes considered jointly and separately in three different analyses (the sample size of these groups permitted estimates to be obtained with the two sexes separately). Individual sub-cohort analyses were meta-analysed.

The estimated contributions of self-reported smoking status (and other covariates) to trait variation in UK Biobank are shown in Supplementary Table 2. These were similar to the ones observed in Generation Scotland, varying between 0.2% (for height) and 1.4% (for waist-to-hip ratio).

Figure 3 shows the proportion of BMI variance explained by the genome-by-smoking interactions in each of the cohorts and sub-cohorts (Generation Scotland, four UK Biobank groups and the UK Biobank meta-analysis). Results for other traits are displayed in Supplementary Figure 2 and full details of the analyses for all traits including estimates, standard errors and log-likelihood ratio tests are shown in Supplementary Tables 4, 5 and 6. Results for the genome-by-smoking-by-sex interactions are shown in Supplementary Figure 3 and Supplementary Table 7.

**Figure 3.**
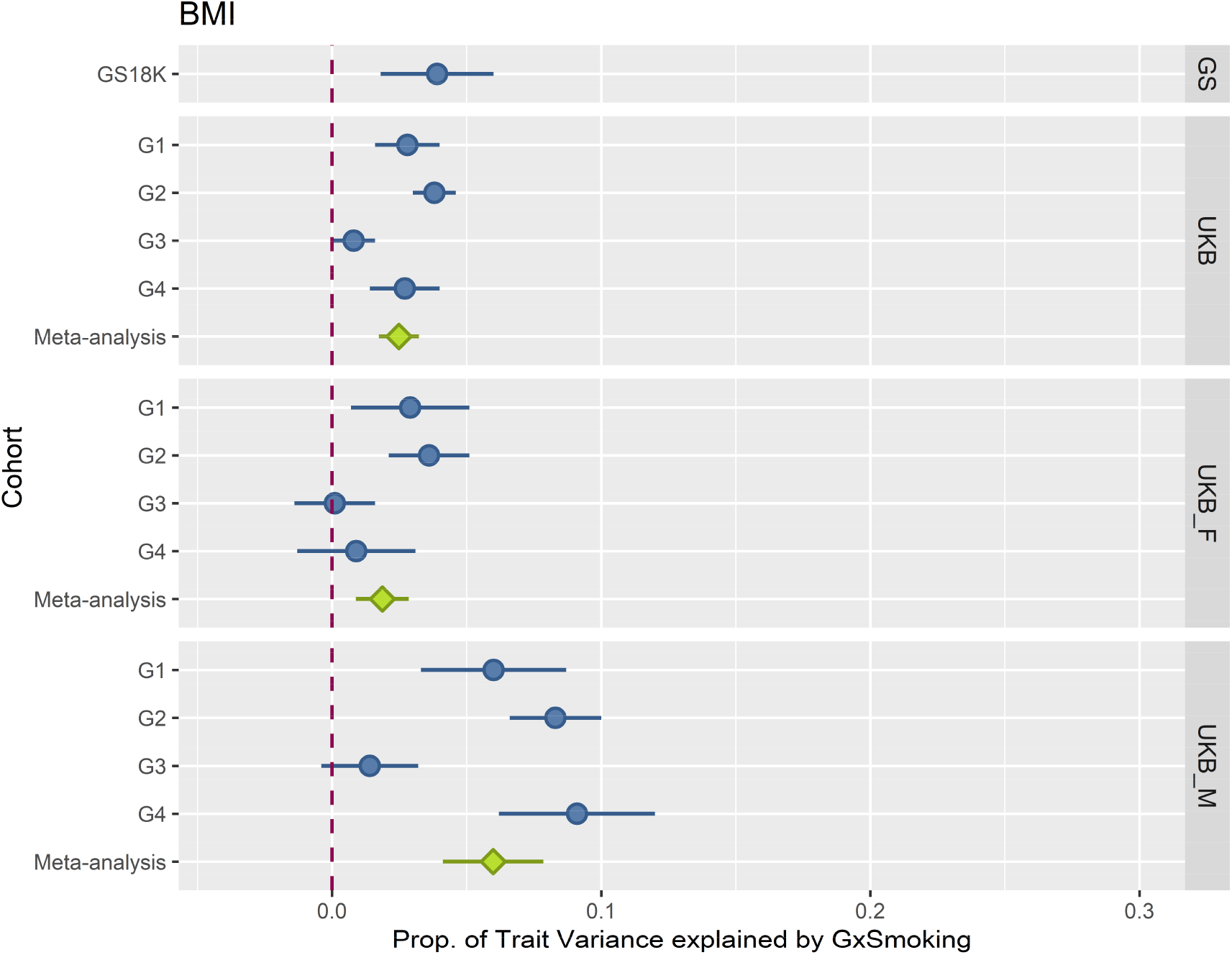
Proportion of BMI variation explained by Genome-by-Smoking interactions across all cohorts and sub-cohorts. The plot shows the proportion of BMI variance (the bars represent standard errors) explained by the genome-by-smoking interaction (x-axis) in the mixed model analyses across cohorts (y-axis). Panels from top to bottom represent cohorts: Generation Scotland (GS), UK Biobank (UKB), UK Biobank females (UKB_F) and UK Biobank males (UKB_M). Blue coloured data points show sub-cohort results, green coloured data points show meta-analyses of the corresponding panel sub-cohorts.

Meta-analyses of the sub-cohorts showed significant genome-by-smoking interactions in all traits except for height when analysing both sexes together and males separately, whereas in females, only fat percentage showed a significant effect of the interaction. Similarly, the genome-by-smoking-by-sex interactions were significant for all traits but height. Genome-by-smoking-by-sex interaction effects explained between 2 and 6% of the observed variation.

### Smoking-associated methylation

To explore the value of DNA methylation data as a proxy for environmental variation, we modelled similarity between individuals based on their DNA methylation levels at a subset of 62 CpG sites previously associated with smoking^19,34^ and which had heritabilities lower than 40%, aiming to target methylation variation that is predominantly capturing environmental variation (for details see Methods). To show that our models can provide accurate estimates we performed a series of simulations. Details and results for those are shown in Supplementary Text 1.

Figure 4 shows the estimates of the proportion of BMI variance explained by different sources included in the mixed linear models in ~9K individuals in Generation Scotland (GS9K - right panel) including models with methylation and genome-by-methylation interactions for models with self-reported smoking status fitted as a fixed effect. Results for other traits are displayed in Supplementary Figure 1 and full details of the analyses for all traits including estimates, standard errors and log-likelihood ratio tests, and results for smoking status fitted as a random effect are shown in Supplementary Table 8. Inclusion of the methylation covariance matrix improved the models for all traits and explained 0.7% of the variance for height and between 3-5% of the variance for obesity-related traits. After including smoking-associated methylation variation, the variation explained by self-reported smoking status dropped to zero for all traits (Supplementary Table 8, Model=GKEM). When exploring the interactions with self-reported smoking status, the estimates in the subset of individuals with methylation data available (N ~ 9K) are substantially larger than in the whole cohort. For example, for BMI, the size of the genome-by-smoking component increased from 4% (GxSmk) to 13% (GxM), however, due to the large standard errors, these two estimates are not significantly different from each other. Inclusion of the genome-by-methylation interaction component nominally improved the model fit for weight, BMI, and waist circumference, with estimates of the interaction component of over 20% of the estimates are large. When fitting jointly the two interaction components (genome-by-smoking and genome-by-methylation) the estimates were not significant for either interaction component (or just nominally significant in the case of genome-by-methylation for BMI). The genome-by-methylation component was also not significant for any trait.

**Figure 4.**
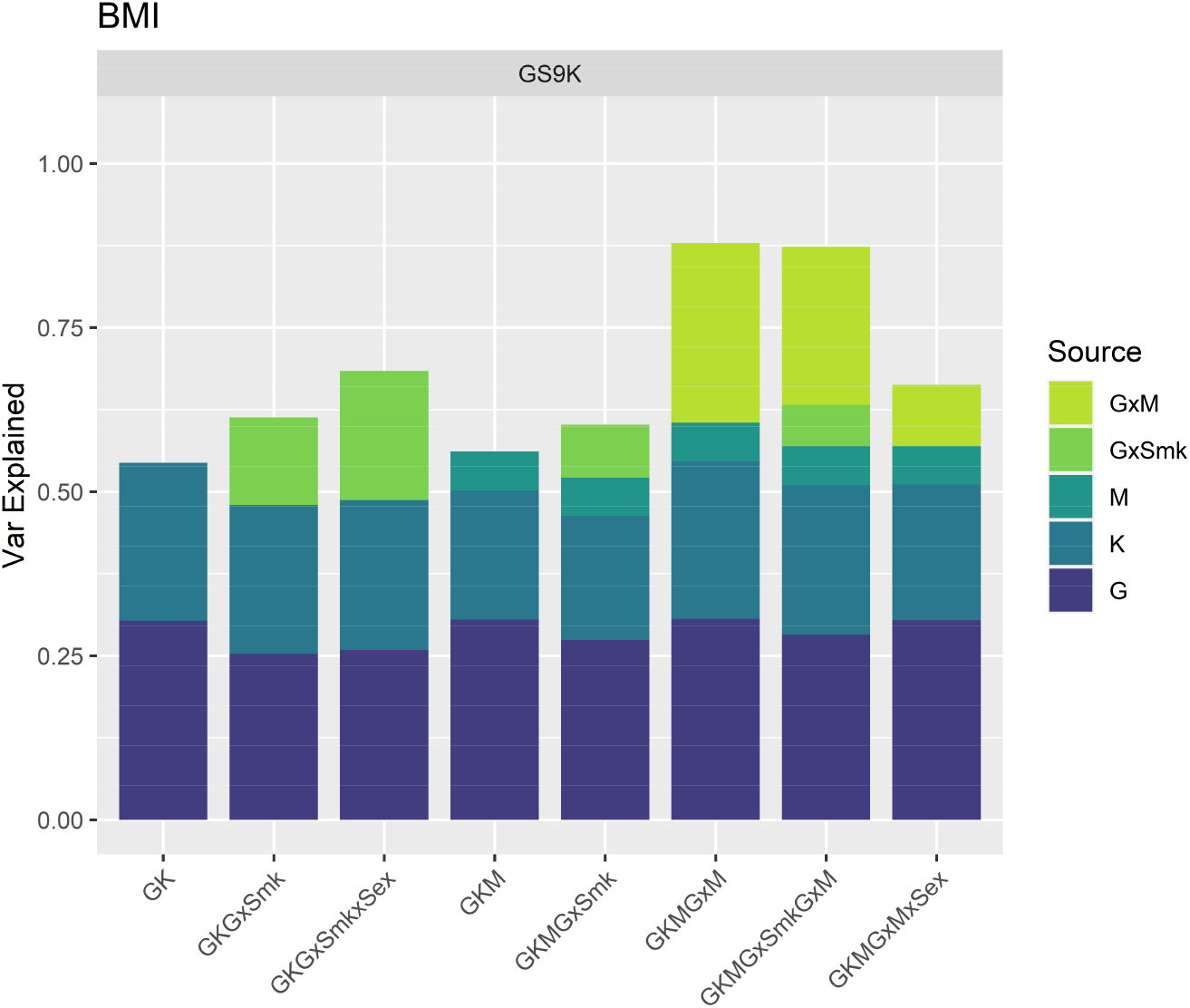
Proportion of BMI variation explained by genetic, environmental and methylation sources in GS9K. Proportion of BMI variance (y-axis) explained by each of the genetic, environmental and interaction sources in the corresponding models (x-axis). G: Genomic, K: Kinship, GxSmk: Genome-by-Smoking, M: Smoking associated methylation, GxM: Genome-by-Methylation, GxSmkxSex: Genome-by-Smoking-by-Sex, GxMxSex: Genome-by-Methylation-by-Sex.

## Discussion

Most complex diseases have moderate heritabilities, with various environmental sources of variation, for example, lifestyle and socioeconomic differences between individuals, also contributing to disease risk^5^. These diseases, particularly obesity, pose major challenges for public health and are associated with heavy economic burdens^3,4,38^. To prevent the problems resulting from complex diseases, effective personalised approaches that help individuals to reach and maintain a healthy lifestyle are required. To achieve that aim, knowledge of environmental effects and gene-by environment interactions (GxE, i.e., understanding the differential effects of an environmental exposure on a trait in individuals with different genotypes^39^) is required. This is a challenge, particularly for environmental factors that are not easy to measure, or that are measured with a lot of error. It has previously been assumed that GxE effects contribute to variation in obesity-related traits^6,8^, but the total contribution to trait variation was not known. Previous analyses exploring GxE in obesity, as well as other traits, took advantage of particular individual genetic variants with known effects, or constructed polygenic scores, combining several genetic variants which reflect genetic risks for the individuals^40,41^. Here we analysed contributions of interactions between the genome (as a whole) with smoking, both using self-reported measures of smoking and methylation data as a proxy for smoking.

Our estimates of the effects of genome-by-smoking interactions in obesity-related traits are larger than those estimated in Shin and Lee ^17^ but in line with Robinson et al. ^12^ for BMI. However, our analyses indicate that the magnitude is substantially different in the two sexes, with interactions playing a bigger role in males for most traits studied (weight, waist, hips, fat%). Joint Analysis of males and females provides less accurate estimates, suggesting that splitting the sexes or modelling the interactions with sex is a more sensible way of analysing the data. The estimates of the variance explained by the interaction components obtained from the genome-by-methylation analyses were large, with also large standard errors. These results, despite not being significant after multiple correction testing, are potentially interesting and should be investigated further. Some studies have suggested that there is potential confounding between interaction and covariance effects in linear mixed models. The CpG sites used to model the methylation similarity between individuals were previously corrected for genomic effects (see methods) removing potential covariance between the genetic and methylation effects ^42,43^.

We estimated that the impact of genome-by-smoking interaction ranges from between 5 to 10% of variation in the studied traits with the exception of height, which we used as a negative control. Our results suggest a larger interaction component in traits associated with weight (BMI, weight, waist, hips) than in those more related to adiposity (waist-to-hip ratio, fat percentage). Biological interpretation of these interactions implies that some genes contributing to obesity differences between individuals have different effects depending on smoking status. This could be mediated in several ways, for example, via genetic variants that affect both obesity and smoking. Some metabolic factors associated with food intake, such as leptin, are suspected to play a role in smoking behaviours, and rewarding effects of food and nicotine are partly mediated by common neurobiological pathways^44^. For example, if these common genetic architectures balance the two behaviours (i.e., more tobacco consumption leading to eating less^44^) the genetic effects of obesity-related traits will be different depending on the smoking status. The interactions could also be driven by gene-by-gene interactions (GxG), i.e., genetic variants affecting obesity modulated by smoking associated genetic variants. Under this scenario smoking status would be capturing smoking associated variants, and the genome-by-smoking interaction would represent GxG instead of GxE. However, given the relatively small heritability of tobacco smoking (SNP heritability ~18%^45^), it is unlikely that all the variation we detected is driven by GxG.

One of the sub-groups of UK Biobank (G3) showed consistently non-significant estimates of the interactions for all traits. The different behaviour for this cohort is not driven by characteristics like the proportion of smokers (Supplementary table 3), or by its genetic stratification. Without any other evidence we cannot attribute these systematic lower estimates to anything but chance.

When we estimated the effect of smoking using the methylomic proxy (62 CpG sites associated with smoking from two independent studies^19,34^), the smoking associated variance increased substantially for all traits (from 2% to 6% for BMI). The methylation component captured the same variance as the self-reported component and some extra variation (Supplementary table 8b). This increase in variation captured could be due to a better ability to separate differences between different levels of smoking (e.g., the self-reported status does not include amount of tobacco smoked, while the methylation might be able to capture this information better). These smoking associated CpG sites could also be picking up variation from other environmental sources that are not exclusively driven by smoking, but correlated with it, such as alcohol intake. When checking in the literature for other possible associations between the 62 CpG sites and other environmental measures (Supplementary Table 9), 20 of these CpGs have previously been associated with age, 15 with alcohol intake or alcohol dependence, 11 with educational attainment, 10 with different types of cancer; and a few with other diseases^46,47^. Unlike for smoking, for most of these associations with other traits, it is unclear if they are casual, or if they could as well be driven by smoking (e.g., alcohol consumption is associated with smoking and picking up a smoking signal).

The fact that variation in obesity can be explained by CpG sites associated with smoking does not imply a causal effect of smoking or methylation on obesity. Methylation is affected by both genetic and environmental effects. Here we selected a subset of CpG sites with moderate to small heritability (lower than 40%, Supplementary Table 9) and we modelled them jointly with a genomic similarity matrix, making it unlikely that the variance picked up by the methylation matrix is genetic in nature. While most changes in methylation at these CpG sites are thought to be causally driven by smoking^19^, associations between methylation and other complex traits, such as BMI, are less well characterised and mostly likely to be reversely caused^48^ (i.e., BMI affecting methylation), however, since our aim was to use methylation as a proxy for the environment, causality does not impact the conclusion of the study. It is, however, important to notice the variable nature of the methylation data, which will change during the life course of individuals unlike the genetics of the individuals, making the inclusion of methylation, measured far back in time, less relevant in a prediction framework^49^. Although this approach should be useful in other populations, a relevant set of CpG sites should be selected reflecting demographic and ethnic relevant associations^50^.

To conclude, we showed that methylation data can be used as a proxy to assess smoking contributions to complex trait variation. We used DNA methylation levels at CpG sites associated with smoking as a proxy for smoking status to assess the contribution of smoking to variation in obesity-related traits. This principle could be extended to take advantage of the wealth of uncovered associations between various *omics* and environmental exposures of interest, particularly for those that are difficult to measure. In humans, relevant interactions could be investigated by exploiting the links between methylation and alcohol intake, metabolomics and diets, the gut microbiome, and diets, etc., and expanding to other species, between the gut microbiome and greenhouse emissions in cattle. This could help expanding our knowledge on their contribution to complex phenotypes, and potentially, help understand the underlying biology and to improve prediction and prognosis.

## Methods

### Data

#### Generation Scotland

We used data from Generation Scotland: Scottish Family Health Study (GS)^51,52^. Ethical approval for the study was given by the NHS Tayside committee on research ethics (ref: 05/s1401/89). Governance of the study, including public engagement, protocol development and access arrangements, was overseen by an independent advisory board, established by the Scottish government. Research participants gave consent to allow both academic and commercial research. Individuals were genotyped with the Illumina HumanOmniExpressExome-8 v1.0 or v1.2. We used PLINK version 1.9b2c^53^ to exclude SNPs that had a missingness > 2% and a Hardy-Weinberg Equilibrium test P < 10^−6^. Markers with a minor allele frequency (MAF) smaller than 0.05 were discarded. Duplicate samples, individuals with gender discrepancies and those with more than 2% missing genotypes were also removed. The resulting data set was merged with the 1092 individuals of the 1000 Genomes population^54^ and a principal component analysis was performed using GCTA^55^. Individuals more than 6 standard deviations away from the mean of principal component 1 and principal component 2 were removed as potentially having African/Asian ancestry as shown in Amador et al.^56^. After quality control, individuals had genotypes for 519,819 common SNP spread over the 22 autosomes. Of the ~24,000 individuals in GS, the number of individuals with complete information for smoking and other covariates was 18,522 so we used this core set of samples for the analyses in order to allow comparisons between the models, we refer to this set of samples as GS18K.

#### UK Biobank

Data access to UK Biobank was granted under MAF 19655. The UK Biobank database include 502,664 participants, aged 40–69, recruited from the general UK population across 22 centres between 2006 and 2010^57^. They underwent extensive phenotyping by questionnaire and clinic measures and provided a blood sample. All participants gave written informed consent, and the study was approved by the North West Multicentre Research Ethics Committee. Phenotypes and genotypes were downloaded direct from UK Biobank. UK Biobank participants were genotyped on two slightly different arrays and quality control was performed by UK Biobank. The two are Affymetrix arrays with 96% of SNPs overlap between both. Further information about the quality control can be found in the UK Biobank website (https://www.ukbiobank.ac.uk/register-apply/). Only genetically white British individuals were used in the analyses. The total number of individuals with complete information for measures of interest was 374,453. Genotypes were available for 534,427 common markers spread over the 22 autosomes.

For computational reasons, UKB individuals were split in four sub-cohorts to be analysed separately. The grouping was based in latitudinal differences between the assessment centres the individuals attended. Number of individuals and assessment centres are shown in Supplementary Table 3.

### Phenotypes

#### Generation Scotland

We used measured phenotypes for eight traits: height, weight, body mass index (BMI, computed as weight/height^2^), waist circumference (waist), hip circumference (hips), waist-to-hip ratio (WHR, computed as waist/hips), bio-impedance analysis fat (fat%), and HDL cholesterol. Phenotypes with values greater or smaller than the mean ± 4 standard deviations (after transformation and adjusting for sex, age and age^2^) were set to missing. The traits were pre-adjusted for the effects of sex, age, age^2^, clinic where the measures were taken, and a rank-based inverse normal transformation was performed on the residuals. These values were used in all the analyses.

#### UK Biobank

We used measured phenotypes for anthropometric traits: height, weight, body mass index (BMI, computed as weight/height^2^),waist circumference (waist), hip circumference (hips), waist-to-hip ratio (WHR, computed as waist/hips), body fat percentage (fat%)Phenotypes with values greater or smaller than the mean ± 4 standard deviations (after transformation and adjusting for sex, age and age^2^) were set to missing. The traits were pre-adjusted for the effects of sex, age, age^2^, clinic where the measures were taken, and a rank-based inverse normal transformation was performed on the residuals. These values were used in all the analyses.

### Smoking status

We used self-reported smoking status on both cohorts. Individuals were classified with respect of smoking as “never smoked”, “ex-smoker” and “current smoker” for Generation Scotland, and as “never smoked”, “ex-smoker”, “current smoker”, and “occasional smoker” for UK Biobank. The number of individuals in each category are shown in Supplementary Table 3.

### DNA Methylation data

DNA methylation data is available for a subset of 9,537 participants from the GS cohort, as part of the Stratifying Resilience and Depression Longitudinally (STRADL) project^58^. From those, we used N = 8,821 individuals that had complete information for all the same set of covariates as used in the smoking status analysis. We refer to this subset of individuals as GS9K. DNA methylation was measured at 866,836 CpGs from whole blood genomic DNA, using the Illumina Infinium MethylationEPIC array. Quality control was performed using R (version 3.6.0)^59^, and packages *shinyMethyl^60^* and *meffil^61^*. We removed outliers based on overall array signal intensity and control probe performance and samples showing a mismatch between recorded and predicted sex. We removed samples with more than 0.5% of sites with a detection p-value of > 0.01; and probes with more than 5% samples with a bead count smaller than 3. Normalization was performed using the R package *minfi^62^*, that produced methylation M-values that were used in downstream analyses. For each methylation site, two linear mixed model were used to remove effects of technical and biological factors correcting for technical variation, i.e., Sentrix id, Sentrix position, batch, clinic, appointment date, year and weekday of the blood extraction, and 20 principal components of the control probes; and biological variation, i.e., sex, age, estimated cell proportions (CD8T, CD4T, NK, B Cell, Mono, and Gran cells proportions based on Houseman, et al. ^63^), and two genetic (Genetic and Kinship) and three common environment (Family, Couples, Siblings) effects. For more information see Xia et al^37^ and Zeng et al^18^. The residual values of those corrections were used for subsequent analyses.

#### Smoking associated CpG sites

We selected a subset of CpG sites identified in two epigenome-wide association studies of tobacco consumption^19,34^. We selected CpG sites with a p-value lower than 10^-7^ in both Ambatipudi et al.^19^ (associations between CpG sites and differences between groups: smokers *v* non-smokers, smokers *v* ex-smokers, ex-smokers *v* non-smokers) and in Joehanes et al.^34^ (associations between CpG sites and dosage of tobacco smoked) to obtain a subset of CpG sites confidently associated with smoking (i.e., from two sources). We identified those CpG sites with heritabilities lower than 40% in Generation Scotland (as measured in the last step of the quality control of the data, see below) that are available in Generation Scotland. The list of 62 CpG sites is available in Supplementary Table 9.

### Covariance Matrices

To model the different sources of variance we used a set of covariance matrices representing similarity between individuals based on genetic components, environmental components, or both.

#### Genetic matrices

**G** is a genomic relationship matrix (GRM) reflecting the genetic similarity between individuals^16,64^. **K** is a matrix representing pedigree relationships as in Zaitlen et al.^65^. It is a modification of G obtained by setting those entries in **G** lower than 0.025 to 0.

#### Smoking matrices

**SMK** is a matrix representing common environmental effects shared between individuals with same smoking status i.e., **SMK** contains a value of 1 between individuals in the same smoking category and a 0 between individuals in different categories.

#### Gene-Environment interaction matrices

**GxSmk** is a matrix representing genome-by-smoking interactions. It was computed as the cell-by-cell product (Hadamard or Schur product) of the corresponding **G** and **SMK** matrices. For an element of the **GxSmk** matrix, if the corresponding **G** or the **SMK** elements are close to zero, the **GxSmk** term will be zero or close to zero as well. Therefore, similarity between individuals due to the interactions represented in the **GxSmk** matrices requires similarity at both genetic and environmental level. This method resembles a reaction norm modelling approach^66^.

#### Methylation-derived matrices

**M** is a matrix representing similarity between individuals based on DNA methylation levels at 62 smoking associated CpG sites (see *Smoking associated CpG sites* above). A similarity matrix was created using OSCA v 0.45^67^ using algorithm 3 (i.e., iteratively standardizing probes and individuals). **GxM** is a genome-by-smoking interaction matrix computed as a Hadamard product of **G** and **M**.

### Analyses

We performed several variance component analyses using GCTA^55^, based in the following linear mixed models:

1. *y* = Xβ + *g*_*g*_ + *g*_*kin*_ + *ɛ*
2. *y* = Xβ + *g*_*g*_ + *g*_*kin*_ + *w*_*L*_ + *ɛ*
3. *y* = Xβ + *g*_*g*_ + *g*_*kin*_ + *w*_*L*_ + *gw* + *ɛ*
4. *y* = Xβ + *g*_*g*_ + *g*_*kin*_ + *gw* + *ɛ*

where *y* is an *n* × 1 vector of observed phenotypes with *n* being the number of individuals, β is a vector of fixed effects and **X** is its design matrix, *g*_*g*_ is an *n* × 1 vector of the total additive genetic effects of the individuals captured by genotyped SNPs with *g*_*g*_ ~ *N*(0, **G***σ^2^*_*g*_); *g*_*kin*_ is an *n* × 1 vector of the extra genetic effects associated with the pedigree for relatives with *g*_*kin*_ ~ *N*(0, K*σ^2^*_*k*_). *w* is a *n* × 1 vector representing the common environmental effects of smoking, with *w* ~ *N*(0, **SMK***σ^2^*_*w*_). *gw* is a *n* × 1 vector representing interactions between markers and environments with *gw* ~ *N*(0, **GxSmk***σ^2^*_*gw*_). *ε* is an *n* × 1 vector for the residuals. The four basic models shown above were expanded to include all combinations of random and fixed effects showed in Figure 1.

The estimates for variance explained by the genome-by-smoking components in the four sub-cohorts of UK Biobank were meta-analysed using the R^59^ package *metafor^68^*.

## Data availability

Generation Scotland data are available from the MRC IGC Institutional Data Access / Ethics Committee for researchers who meet the criteria for access to confidential data. Generation Scotland data are available to researchers on application to the Generation Scotland Access Committee. The managed access process ensures that approval is granted only to research which comes under the terms of participant consent which does not allow making participant information publicly available. UK Biobank data are available from: https://www.ukbiobank.ac.uk/register-apply/

## Acknowledgements

The authors want to acknowledge the Medical Research Council (MRC) UK for funding (core funding: MC_UU_00007/10, MC_PC_U127592696 and MC_PC_U127561128), and a *Wellcome Trust Investigator Award* to AMM (220857/Z/20/Z). Generation Scotland has received core funding from the Chief Scientist Office of the Scottish Government Health Directorates CZD/16/6 and the Scottish Funding Council HR03006. Genotyping of the GS samples was carried out by the Genetics Core Laboratory at the Wellcome Trust Clinical Research Facility, Edinburgh, Scotland and was funded by the UK MRC and the Wellcome Trust (Wellcome Trust Strategic Award “STratifying Resilience and Depression Longitudinally” (STRADL) Reference 104036/Z/14/Z). We are grateful to all the families who took part, the general practitioners, and the Scottish School of Primary Care for their help in recruiting them, and the whole Generation Scotland team, which includes interviewers, computer and laboratory technicians, clerical workers, research scientists, volunteers, managers, receptionists, healthcare assistants and nurses. The research was conducted using the UK Biobank Resource under Application number 19655.

## Author Contributions

CA, CHS, and PN conceived and designed the experiments presented in this manuscript. DP and AMM contributed to conceive and design the study population and phenotypic recording. DP, AMM, and JFW contributed to oversight of the study and sample collection. CA conducted the analyses. YZ, MB, RW, KE, AC, CH, managed and maintained the data and performed quality control and data annotation. CA, PN, CSH wrote the paper. All authors discussed results, read, and approved the final manuscript.

## Supplementary Figure and Table captions

**Supplementary Figure 1. Proportion of trait variation explained by the different sources in Generation Scotland (GS) in each of the eight traits studied**. Proportion of trait variance (y-axis) explained by each of the genetic, environmental and interaction sources in the corresponding models (x-axis). Left panel: GS data (Nind≈18K) with complete environmental information. Right panel: GS data with methylation information (Nind≈9K). G: Genomic, K: Kinship, GxSmk: Genome-by-Smoking, M: Smoking associated methylation, GxM: Genome-by-Methylation, GxSmkxSex: Genome-by-Smoking-by-Sex, GxMxSex: Genome-by-Methylation-by-Sex.

**Supplementary Figure 2. Proportion of trait variation explained by Genome-by-Smoking interactions across all cohorts and sub-cohorts in each of the eight traits studied**. The plot shows the proportion of trait variance (the bars represent standard errors) explained by the genome-by-smoking interaction (x-axis) in the mixed model analyses across cohorts (y-axis). Panels from top to bottom represent cohorts: Generation Scotland (GS), UK Biobank (UKB), UK Biobank females (UKB_F) and UK Biobank males (UKB_M). Blue coloured data points show sub-cohort results (GS18K and UKB subgroups G1-G4), green coloured data points show meta-analyses of the corresponding panel sub-cohorts.

**Supplementary Figure 3. Proportion of trait variation explained by Genome-by-Smoking-by-Sex interactions across all cohorts and sub-cohorts in each of the eight traits studied**. The plot shows the proportion of BMI variance (the bars represent standard errors) explained by the genome-by-smoking-by-sex interaction (x-axis) in the mixed model analyses across cohorts (y-axis). Panels from top to bottom represent cohorts: Generation Scotland (GS), UK Biobank (UKB), UK Biobank females (UKB_F) and UK Biobank males (UKB_M). Blue coloured data points show sub-cohort results (GS18K and UKB subgroups G1-G4), green coloured data points show meta-analyses of the corresponding panel sub-cohorts.

**Supplementary Table 1. Results for all models for GS18K cohort**. A. Models with smoking fitted as a random effect. B. Models with smoking fitted as a random effect. The tables show, for each trait, proportion of the phenotypic variance explained (Var), standard error (SE), Significance of the t-statistic (Sig, P), P value for the log-likelihood ratio test (LRT P, only for the interactions) by each of the components in the model: Genetic (G), Kinship (K), Smoking (when fitted as a random effect, Smk), genome-by-smoking interaction (GxSmk), genome-by-smoking-by-sex interaction (GxSmkxSex), kinship-by-smoking interaction (KxSmk). Highlighted P values indicate nominally significant results for the interaction components.

**Supplementary Table 2. Variance explained by fixed effects**. Percentage of the phenotypic variance explained by the fixed effects included in the models for each trait and cohort.

**Supplementary Table 3. Cohorts summaries**. Summary statistics (number of individuals in each category or mean values) for the covariates included in the models for each of the analysed cohorts.

**Supplementary Table 4. Results for all models for the four UKB cohorts (joint sexes)**. The tables show, for each trait, proportion of the phenotypic variance explained (Var), standard error (SE), Significance of the t-statistic (Sig, P), P value for the log-likelihood ratio test (LRT P, only for the interactions) by each of the components in the model: Genetic (G), Kinship (K), genome-by-smoking interaction (GxSmk). Highlighted P values indicate nominally significant results for the interaction components in each of the four sub-cohorts of UK Biobank (G1, G2, G3, G4) and their Meta-Analyses.

**Supplementary Table 5. Results for all models for the four UKB cohorts (males)**. The tables show, for each trait, proportion of the phenotypic variance explained (Var), standard error (SE), Significance of the t-statistic (Sig, P), P value for the log-likelihood ratio test (LRT P, only for the interactions) by each of the components in the model: Genetic (G), Kinship (K), genome-by-smoking interaction (GxSmk). Highlighted P values indicate nominally significant results for the interaction components in males from each of the four sub-cohorts of UK Biobank (G1_M, G2_M, G3_M, G4_M) and their Meta-Analyses.

**Supplementary Table 6. Results for all models for the four UKB cohorts (females)**. The tables show, for each trait, proportion of the phenotypic variance explained (Var), standard error (SE), Significance of the t-statistic (Sig, P), P value for the log-likelihood ratio test (LRT P, only for the interactions) by each of the components in the model: Genetic (G), Kinship (K), genome-by-smoking interaction (GxSmk). Highlighted P values indicate nominally significant results for the interaction components in females from each of the four sub-cohorts of UK Biobank (G1_F, G2_F, G3_F, G4_F) and their Meta-Analyses.

**Supplementary Table 7. Results for all models for the four UKB cohorts (joint GxSmkxSex interactions)**. The tables show, for each trait, proportion of the phenotypic variance explained (Var), standard error (SE), Significance of the t-statistic (Sig, P), P value for the log-likelihood ratio test (LRT P, only for the interactions) by each of the components in the model: Genetic (G), Kinship (K), genome-by-smoking-by-sex interaction (GxSmkxSex). Highlighted P values indicate nominally significant results for the interaction components in each of the four sub-cohorts of UK Biobank (G1, G2, G3, G4) and their Meta-Analyses.

**Supplementary Table 8. Results for all models for GS9K cohort**. A. Models with smoking fitted as a random effect. B. Models with smoking fitted as a random effect. The tables show, for each trait, proportion of the phenotypic variance explained (Var), standard error (SE), Significance of the t-statistic (Sig, P), P value for the log-likelihood ratio test (LRT P, only for the interactions) by each of the components in the model: Genetic (G), Kinship (K), Smoking (when fitted as a random effect, Smk), genome-by-smoking interaction (GxSmk), genome-by-smoking-by-sex interaction (GxSmkxSex), kinship-by-smoking interaction (KxSmk). Highlighted P values indicate nominally significant results for the interaction components.

**Supplementary Table 9. Smoking associated CpG sites information**. Name, chromosome, location, heritability, and trait associations of the 62 CpG sites associated with smoking. Trait associations were extracted from the EWAS Atlas database.

## References

1. Locke, A.E. et al. Genetic studies of body mass index yield new insights for obesity biology. Nature 518, 197–206 (2015).

2. Qi, L. & Cho, Y.A. Gene-environment interaction and obesity. Nutr. Rev. 66, 684–694 (2008).

3. Swinburn, B.A. et al. The global obesity pandemic: shaped by global drivers and local environments. Lancet 378, 804–814 (2011).

4. Wang, Y.C., McPherson, K., Marsh, T., Gortmaker, S.L. & Brown, M. Health and economic burden of the projected obesity trends in the USA and the UK. Lancet 378, 815–825 (2011).

5. Amador, C. et al. Regional variation in health is predominantly driven by lifestyle rather than genetics. Nat. Commun. 8, 801 (2017).

6. Huang, T. & Hu, F.B. Gene-environment interactions and obesity: recent developments and future directions. BMC Med. Genomics. 8, S2 (2015).

7. Tyrrell, J. et al. Gene–obesogenic environment interactions in the UK Biobank study. Int. J. Epidemiol. 46, 559–575 (2017).

8. Cornelis, M.C. & Hu, F.B. Gene-Environment Interactions in the Development of Type 2 Diabetes: Recent Progress and Continuing Challenges. Annu. Rev. Nutr. 32, 245–259 (2012).

9. Li, J., Li, X., Zhang, S. & Snyder, M. Gene-Environment Interaction in the Era of Precision Medicine. Cell 177, 38–44 (2019).

10. Poveda, A. et al. The heritable basis of gene–environment interactions in cardiometabolic traits. Diabetologia 60, 442–452 (2017).

11. Trzaskowski, M., Lichtenstein, P., Magnusson, P.K., Pedersen, N.L. & Plomin, R. Application of linear mixed models to study genetic stability of height and body mass index across countries and time. Int. J. Epidemiol. 45, 417–423 (2016).

12. Robinson, M.R. et al. Genotype-covariate interaction effects and the heritability of adult body mass index. Nat. Genet. 49, 1174–1181 (2017).

13. Bentley, A.R. et al. Multi-ancestry genome-wide gene–smoking interaction study of 387,272 individuals identifies new loci associated with serum lipids. Nat. Genet. 51, 636–648 (2019).

14. Justice, A.E. et al. Genome-wide meta-analysis of 241,258 adults accounting for smoking behaviour identifies novel loci for obesity traits. Nat. Commun. 8, 14977 (2017).

15. Sung, Y.J. et al. A multi-ancestry genome-wide study incorporating gene–smoking interactions identifies multiple new loci for pulse pressure and mean arterial pressure. Hum. Mol. Genet. 28, 2615–2633 (2019).

16. Yang, J. et al. Common SNPs explain a large proportion of the heritability for human height. Nat. Genet. 42, 565–569 (2010).

17. Shin, J. & Lee, S.H. GxEsum: genotype-by-environment interaction model based on summary statistics. bioRxiv (2020).

18. Zeng, Y. et al. Parent of origin genetic effects on methylation in humans are common and influence complex trait variation. Nat. Commun. 10, 1383 (2019).

19. Ambatipudi, S. et al. Tobacco smoking-associated genome-wide DNA methylation changes in the EPIC study. Epigenomics 8, 599–618 (2016).

20. Sugden, K. et al. Establishing a generalized polyepigenetic biomarker for tobacco smoking. Transl. Psychiatry 9, 92 (2019).

21. Zhang, Y. & Kutateladze, T.G. Diet and the epigenome. Nat. Commun. 9, 3375 (2018).

22. McRae, A.F. et al. Contribution of genetic variation to transgenerational inheritance of DNA methylation. Genome Biol. 15, R73 (2014).

23. van Dongen, J. et al. Genetic and environmental influences interact with age and sex in shaping the human methylome. Nat. Commun. 7, 11115 (2016).

24. Jones, P.A. Functions of DNA methylation: islands, start sites, gene bodies and beyond. Nat. Rev. Genet. 13, 484–492 (2012).

25. Moore, L.D., Le, T. & Fan, G. DNA Methylation and Its Basic Function. Neuropsychopharmacology 38, 23–38 (2013).

26. Putiri, E.L. & Robertson, K.D. Epigenetic mechanisms and genome stability. Clin. Epigenetics 2, 299–314 (2011).

27. Greenberg, M.V.C. & Bourc’his, D. The diverse roles of DNA methylation in mammalian development and disease. Nat. Rev. Mol. Cell Biol. 20, 590–607 (2019).

28. Demerath, E.W. et al. Epigenome-wide association study (EWAS) of BMI, BMI change and waist circumference in African American adults identifies multiple replicated loci. Hum. Mol. Genet. 24, 4464–4479 (2015).

29. Wahl, S. et al. Epigenome-wide association study of body mass index, and the adverse outcomes of adiposity. Nature 541, 81 (2016).

30. Rask-Andersen, M. et al. Epigenome-wide association study reveals differential DNA methylation in individuals with a history of myocardial infarction. Hum. Mol. Genet. 25, 4739–4748 (2016).

31. Jin, Z. & Liu, Y. DNA methylation in human diseases. Genes Dis. 5, 1–8 (2018).

32. Horvath, S. DNA methylation age of human tissues and cell types. Genome Biol. 14, 3156 (2013).

33. Horvath, S. & Raj, K. DNA methylation-based biomarkers and the epigenetic clock theory of ageing. Nat. Rev. Genet. 19, 371–384 (2018).

34. Joehanes, R. et al. Epigenetic Signatures of Cigarette Smoking. Circ. Cardiovasc. Genet. 9, 436–447 (2016).

35. Lee, M.K., Hong, Y., Kim, S.-Y., London, S.J. & Kim, W.J. DNA methylation and smoking in Korean adults: epigenome-wide association study. Clin. Epigenetics 8, 103 (2016).

36. Gao, X., Jia, M., Zhang, Y., Breitling, L.P. & Brenner, H. DNA methylation changes of whole blood cells in response to active smoking exposure in adults: a systematic review of DNA methylation studies. Clin. Epigenetics 7, 113 (2015).

37. Xia, C. et al. Pedigree- and SNP-Associated Genetics and Recent Environment are the Major Contributors to Anthropometric and Cardiometabolic Trait Variation. PLOS Genet. 12, e1005804 (2016).

38. Gortmaker, S.L. et al. Changing the future of obesity: science, policy, and action. Lancet 378, 838–847 (2011).

39. Ottman, R. Gene–Environment Interaction: Definitions and Study Design. Prev. Med. 25, 764–770 (1996).

40. Young, A.I., Wauthier, F. & Donnelly, P. Multiple novel gene-by-environment interactions modify the effect of FTO variants on body mass index. Nat. Commun. 7, 12724 (2016).

41. Qi, Q. et al. Sugar-Sweetened Beverages and Genetic Risk of Obesity. N. Engl. J. Med. 367, 1387–1396 (2012).

42. Ni, G. et al. Genotype–covariate correlation and interaction disentangled by a whole-genome multivariate reaction norm model. Nat. Commun. 10, 2239 (2019).

43. Zhou, X., Im, H.K. & Lee, S.H. CORE GREML for estimating covariance between random effects in linear mixed models for complex trait analyses. Nat. Commun. 11, 4208 (2020).

44. Chao, A.M., Wadden, T.A., Ashare, R.L., Loughead, J. & Schmidt, H.D. Tobacco Smoking, Eating Behaviors, and Body Weight: a Review. Curr. Addict. Rep. 6, 191–199 (2019).

45. Evans, L.M. et al. Genetic architecture of four smoking behaviors using partitioned *h*^2^ _*SNP*_. medRxiv, 2020.06.17.20134080 (2020).

46. Battram, T. et al. The EWAS Catalog: a database of epigenome-wide association studies. .https://doi.org/10.31219/osf.io/837wn (2021).

47. Li, M. et al. EWAS Atlas: a curated knowledgebase of epigenome-wide association studies. Nucleic Acids Res 47, D983–d988 (2019).

48. Reed, Z.E., Suderman, M.J., Relton, C.L., Davis, O.S.P. & Hemani, G. The association of DNA methylation with body mass index: distinguishing between predictors and biomarkers. Clin. Epigenetics 12, 50 (2020).

49. Trejo Banos, D. et al. Bayesian reassessment of the epigenetic architecture of complex traits. Nat. Commun. 11, 2865 (2020).

50. Popejoy, A.B. et al. The clinical imperative for inclusivity: Race, ethnicity, and ancestry (REA) in genomics. Hum. Mutat. 39, 1713–1720 (2018).

51. Smith, B.H. et al. Cohort profile: Generation Scotland: Scottish Family Health Study (GS:SFHS). The study, its participants and their potential for genetic research on health and illness. Int. J. Epidemiol. 42, 689–700 (2012).

52. Smith, B.H. et al. Generation Scotland: the Scottish Family Health Study; a new resource for researching genes and heritability. BMC Med. Genet. 7, 74 (2006).

53. Chang, C. et al. Second-generation PLINK: rising to the challenge of larger and richer datasets. GigaScience 4, 7 (2015).

54. The 1000 Genomes Project Consortium. A map of human genome variation from population-scale sequencing. Nature 467, 1061–1073 (2010).

55. Yang, J., Lee, S.H., Goddard, M.E. & Visscher, P.M. GCTA: a tool for genome-wide complex trait analysis. Am. J. Hum. Genet. 88, 76–82 (2011).

56. Amador, C. et al. Recent genomic heritage in Scotland. BMC Genomics 16(2015).

57. Sudlow, C. et al. UK Biobank: An Open Access Resource for Identifying the Causes of a Wide Range of Complex Diseases of Middle and Old Age. PLOS Med. 12, e1001779 (2015).

58. Navrady, L.B. et al. Cohort Profile: Stratifying Resilience and Depression Longitudinally (STRADL): a questionnaire follow-up of Generation Scotland: Scottish Family Health Study (GS:SFHS). Int. J. Epidemiol. 47, 13–14g (2017).

59. R Core Team. R: A language and environment for statistical computing, (R Foundation for Statistical Computing, Vienna (Austria), 2020).

60. Fortin, J., Fertig, E. & Hansen, K. shinyMethyl: interactive quality control of Illumina 450k DNA methylation arrays in R [version 2; peer review: 2 approved]. F1000Research 3(2014).

61. Min, J.L., Hemani, G., Davey Smith, G., Relton, C. & Suderman, M. Meffil: efficient normalization and analysis of very large DNA methylation datasets. Bioinformatics 34, 3983–3989 (2018).

62. Aryee, M.J. et al. Minfi: a flexible and comprehensive Bioconductor package for the analysis of Infinium DNA methylation microarrays. Bioinformatics 30, 1363–1369 (2014).

63. Houseman, E.A. et al. DNA methylation arrays as surrogate measures of cell mixture distribution. BMC Bioinf. 13, 86 (2012).

64. VanRaden, P.M. Efficient methods to compute genomic predictions. J. Dairy Sci. 91, 4414–4423 (2008).

65. Zaitlen, N. et al. Using extended genealogy to estimate components of heritability for 23 quantitative and dichotomous traits. PLOS Genet. 9, e1003520 (2013).

66. Jarquín, D. et al. A reaction norm model for genomic selection using high-dimensional genomic and environmental data. Theor. Appl. Genet. 127, 595–607 (2014).

67. Zhang, F. et al. OSCA: a tool for omic-data-based complex trait analysis. Genome Biol. 20, 107 (2019).

68. Viechtbauer, W. Conducting Meta-Analyses in R with the metafor Package. J Stat Softw 36, 48 (2010).

