## Supplementary Figures (1-3) for "Genome-wide methylation data improves dissection of the effect of smoking on body mass index"

Supplementary Figure 1. Proportion of trait variation explained by the different sources in Generation Scotland (GS) in each of the eight traits studied.

Proportion of trait variance (y-axis) explained by each of the genetic, environmental and interaction sources in the corresponding models (x-axis). Left panel: GS data (Nind≈18K) with complete environmental information. Right panel: GS data with methylation information (Nind≈9K). G: Genomic, K: Kinship, GxSmk: Genome-by-Smoking, M: Smoking associated methylation, GxM: Genome-by-Methylation, GxSmkxSex: Genome-by-Smoking-by-Sex, GxMxSex: Genome-by-Methylation-by-Sex.

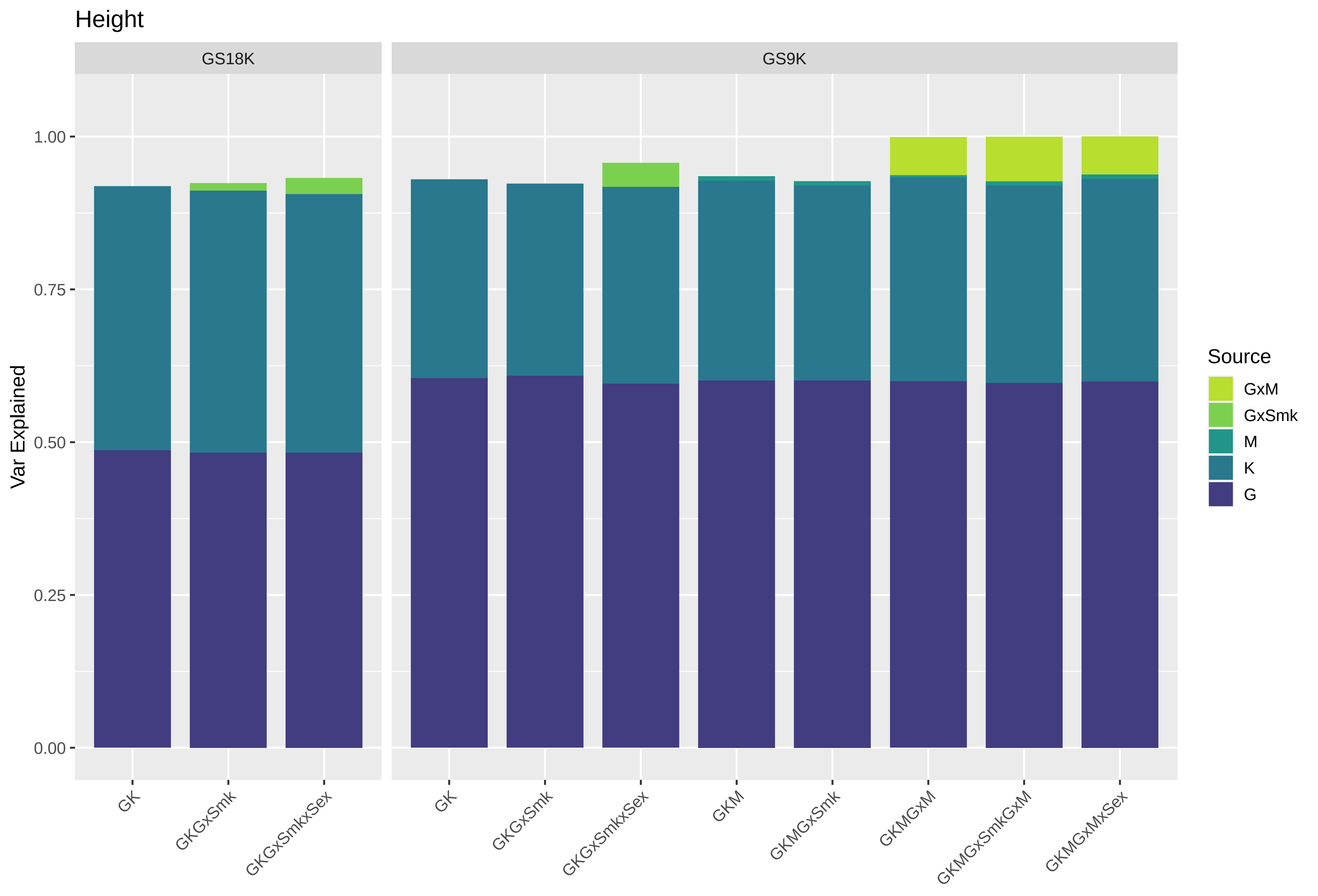

Height
GS18K
GS9K
Var Explained
1.00
0.75
0.50
0.25
0.00
GK
GKGxSmk
GKGxSmkxSex
GK
GKGxSmk
GKGxSmkxSex
GKM
GKMGxSmk
GKMGxM
GKMGxSmkGxM
GKMGxMxSex
Source
GxM
GxSmk
M
K
G

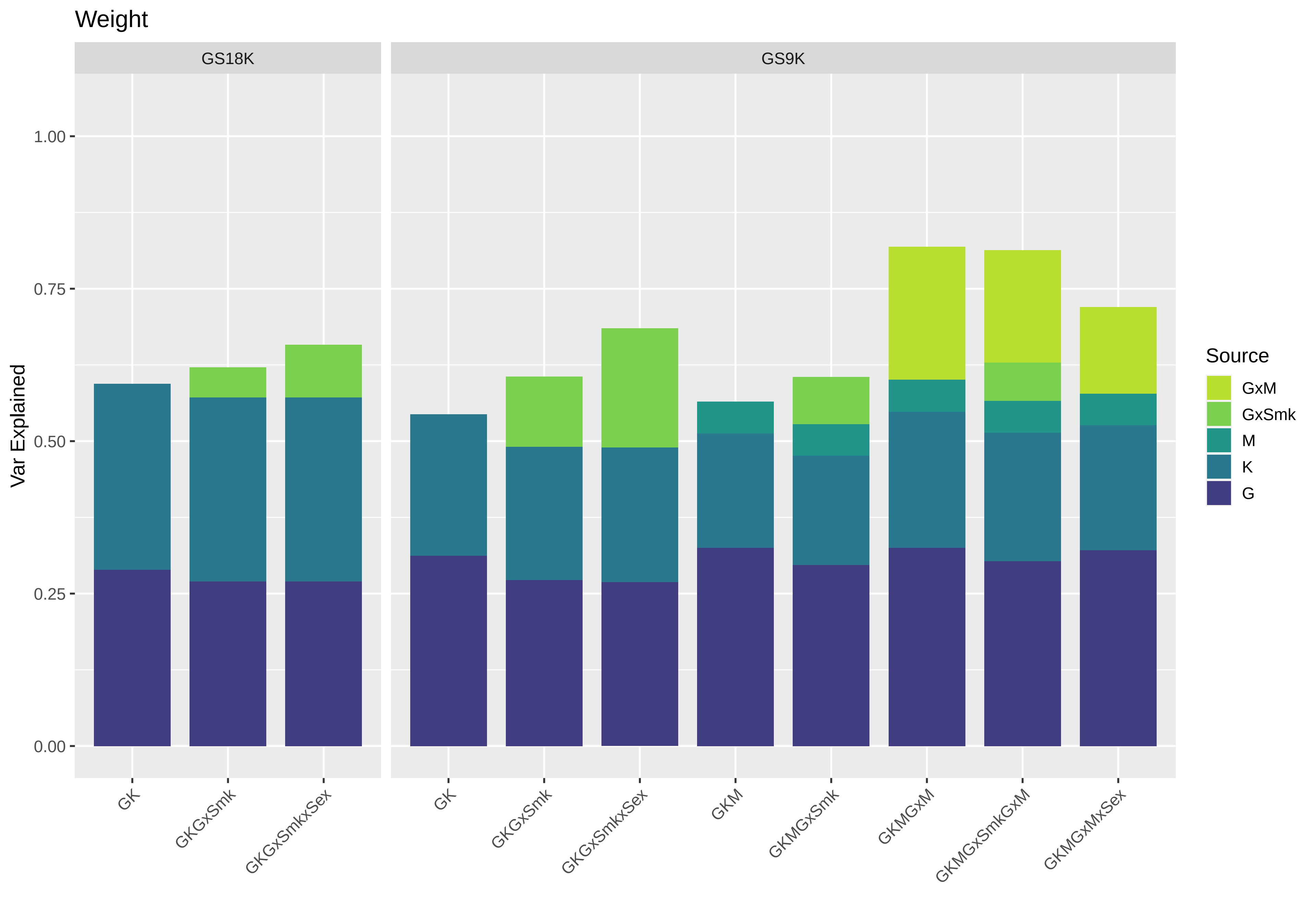
Weight
GS18K
GS9K
Var Explained
1.00
0.75
0.50
0.25
0.00
GK
GKGxSmk
GKGxSmkxSex
GK
GKGxSmk
GKGxSmkxSex
GKM
GKMGxSmk
GKMGxM
GKMGxSmkGxM
GKMGxMxSex
Source
GxM
GxSmk
M
K
G

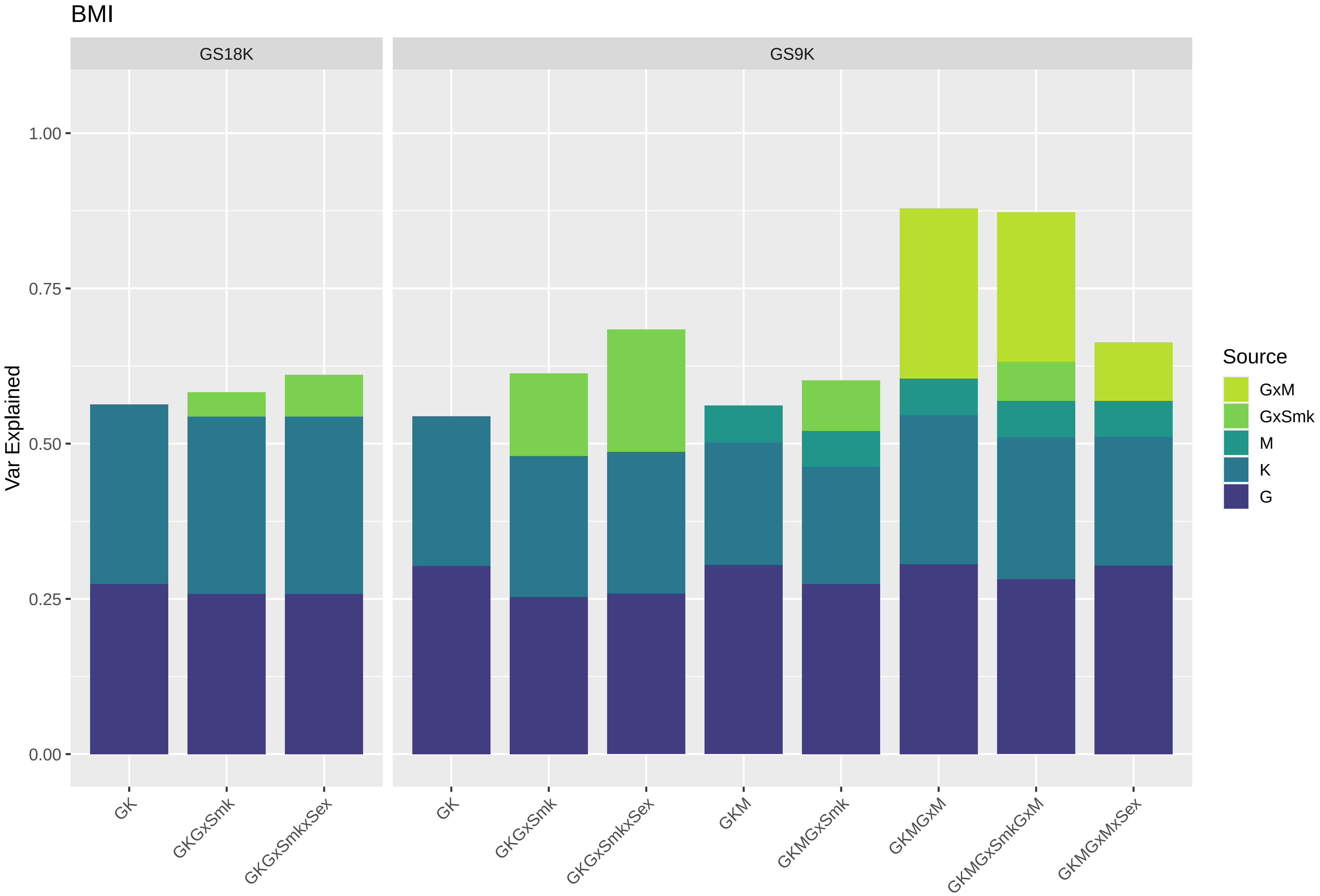
BMI
GS18K
GS9K
Var Explained
0.00
0.25
0.50
0.75
1.00
GK
GKGxSmk
GKGxSmkxSex
GK
GKGxSmk
GKGxSmkxSex
GKM
GKMGxSmk
GKMGxM
GKMGxSmkGxM
GKMGxMxSex
Source
GxM
GxSmk
M
K
G

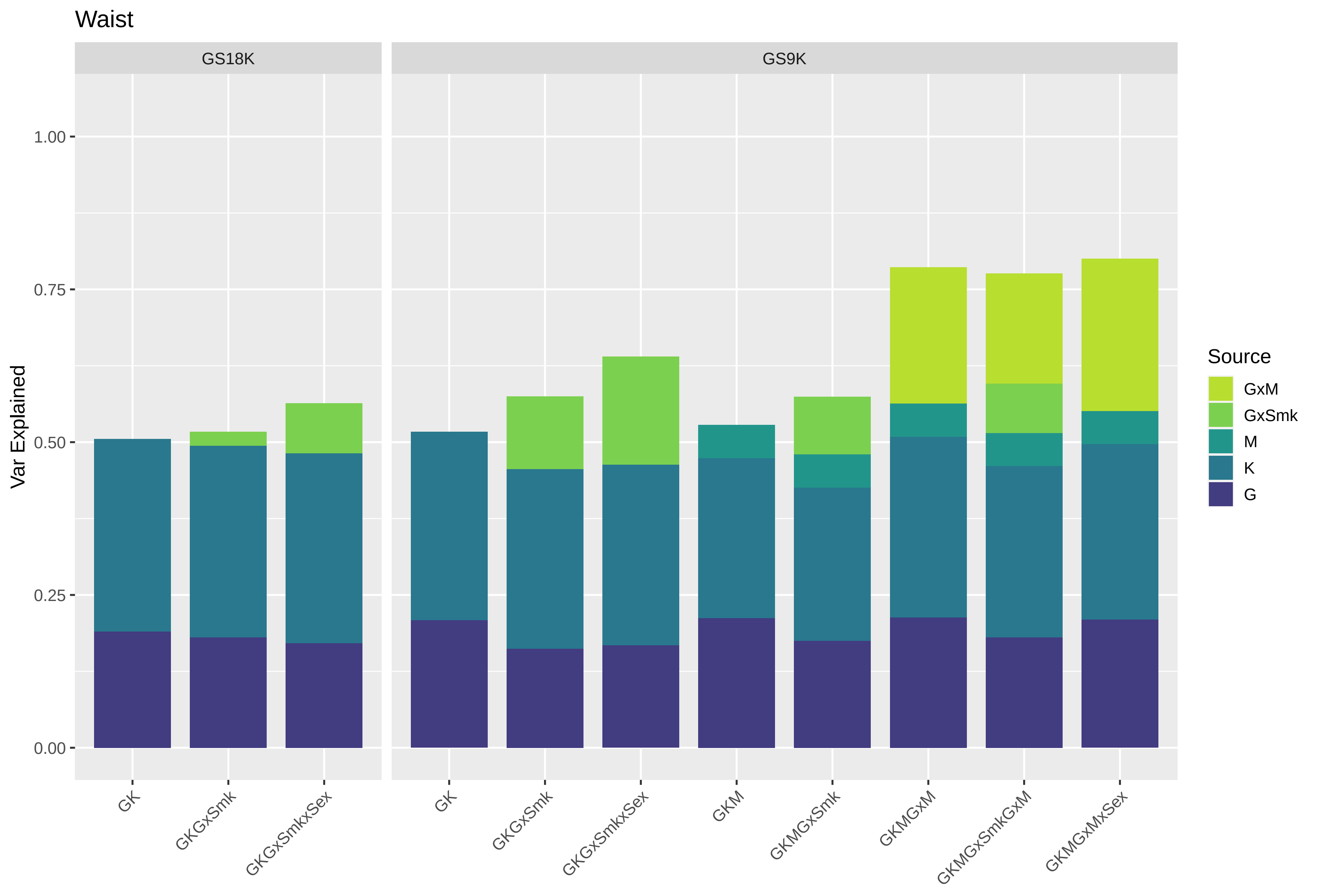
Waist
GS18K
GS9K
Var Explained
1.00
0.75
0.50
0.25
0.00
GK
GKGxSmk
GKGxSmkxSex
GK
GKGxSmk
GKGxSmkxSex
GKM
GKMGxSmk
GKMGxM
GKMGxSmkGxM
GKMGxMxSex
Source
GxM
GxSmk
M
K
G

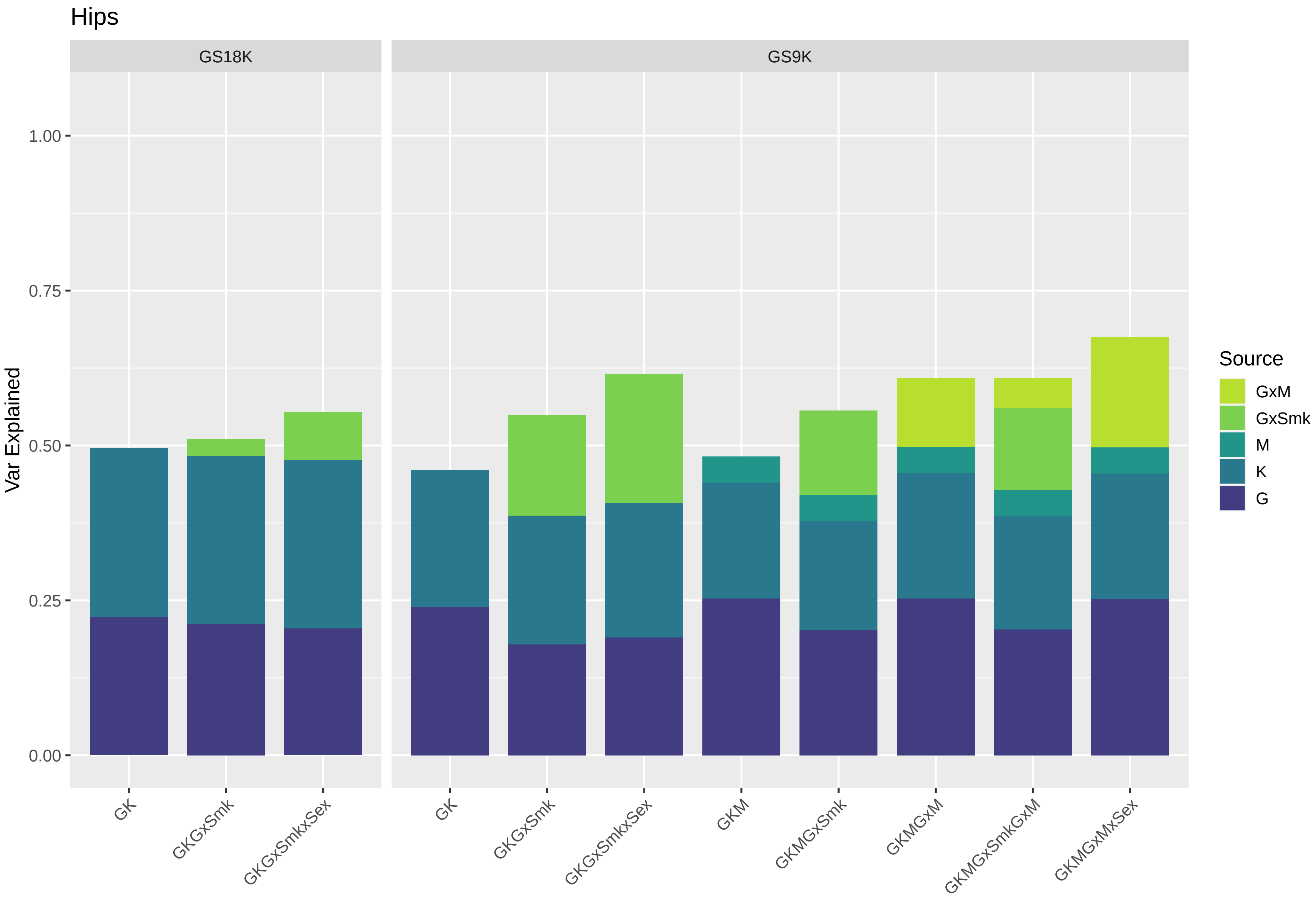

Hips
GS18K
GS9K
Var Explained
1.00
0.75
0.50
0.25
0.00
GK
GKGxSmk
GKGxSmkxSex
GK
GKGxSmk
GKGxSmkxSex
GKM
GKMGxSmk
GKMGxM
GKMGxSmkGxM
GKMGxMxSex
Source
GxM
GxSmk
M
K
G

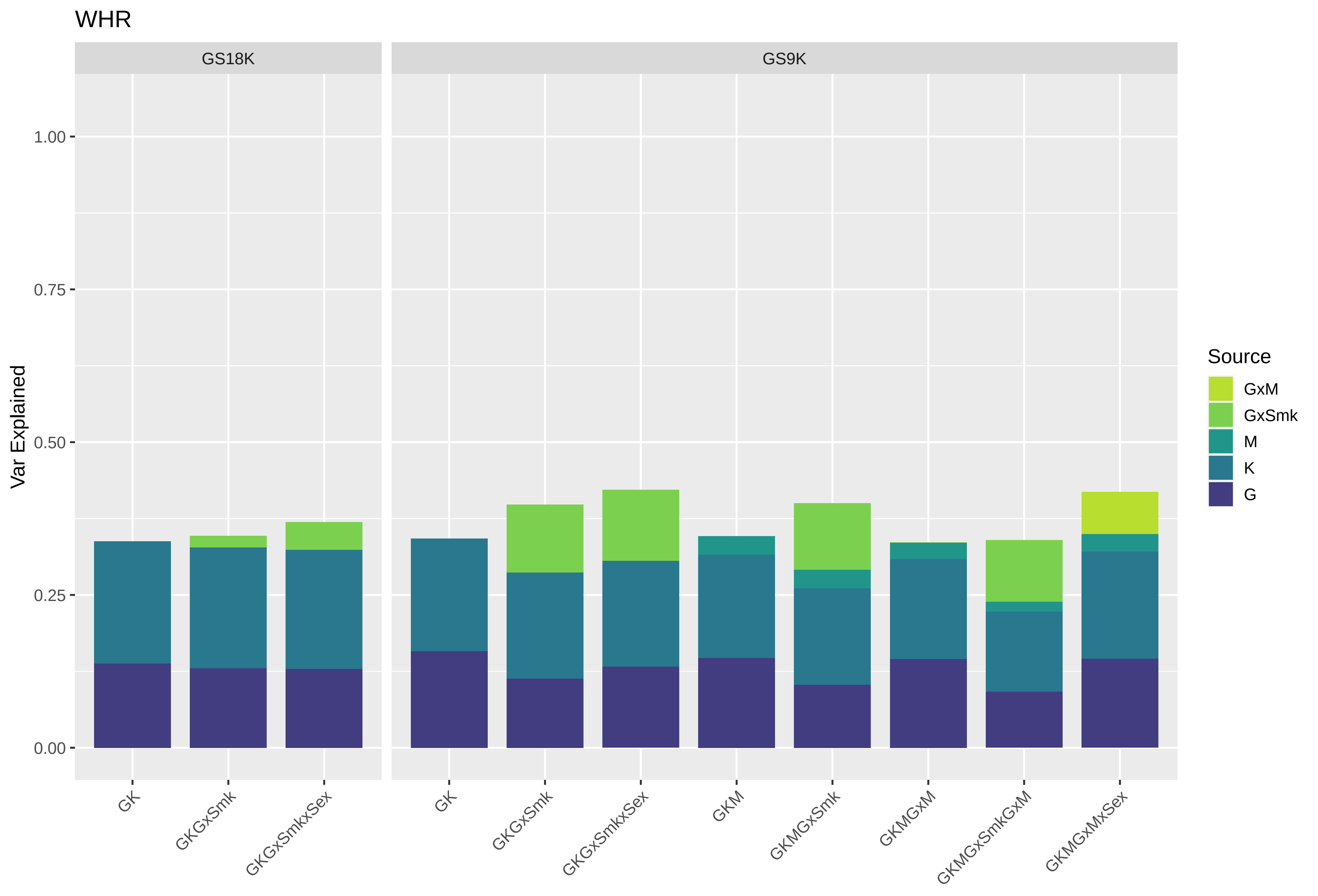

WHR
GS18K
GS9K
Var Explained
1.00
0.75
0.50
0.25
0.00
GK
GKGxSmk
GKGxSmkxSex
GK
GKGxSmk
GKGxSmkxSex
GKM
GKMGxSmk
GKMGxM
GKMGxSmkGxM
GKMGxMxSex
Source
GxM
GxSmk
M
K
G

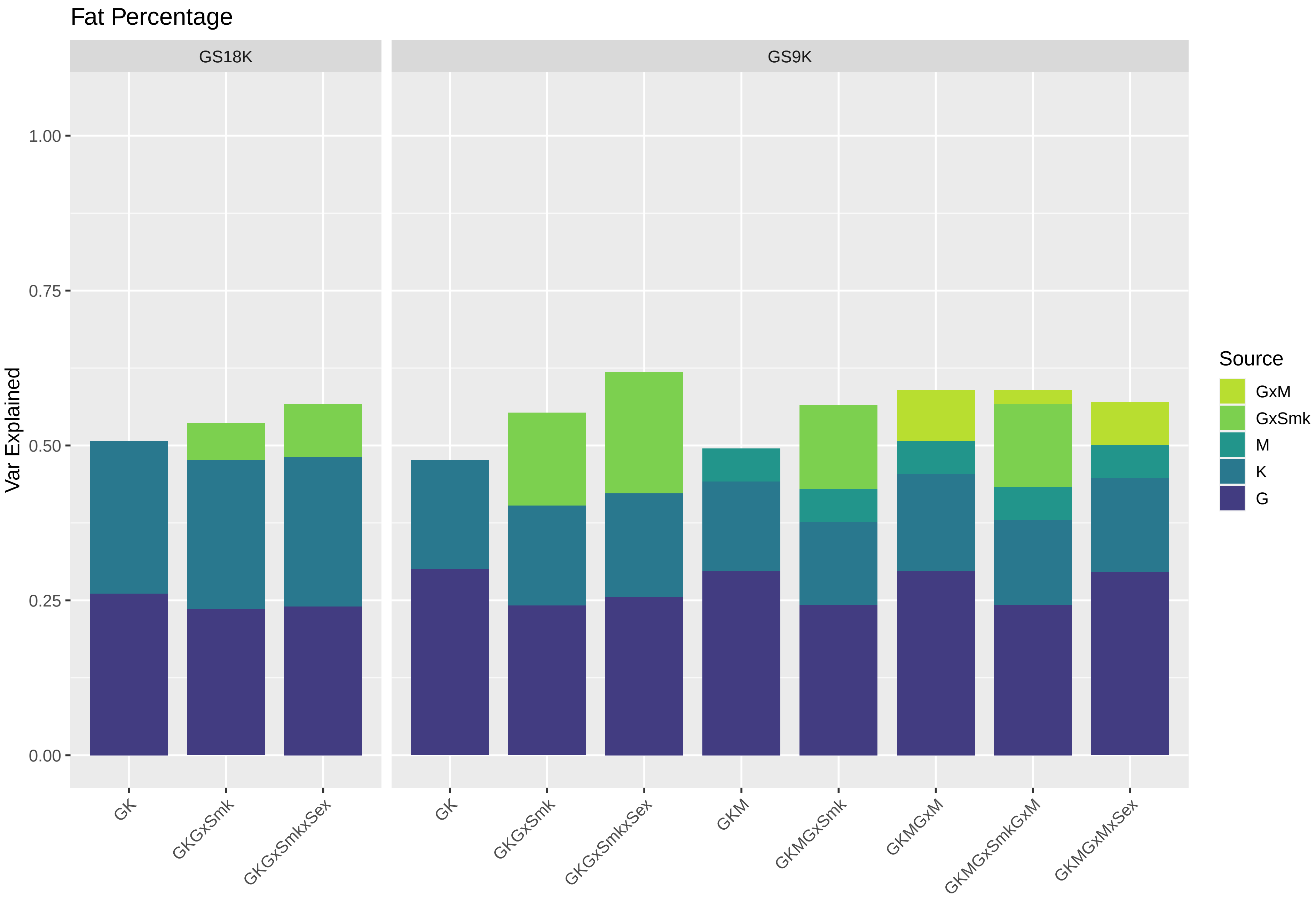

Fat Percentage
GS18K
GS9K
Var Explained
1.00
0.75
0.50
0.25
0.00
GK
GKGxSmk
GKGxSmkxSex
GK
GKGxSmk
GKGxSmkxSex
GKM
GKMGxSmk
GKMGxM
GKMGxSmkGxM
GKMGxMxSex
Source
GxM
GxSmk
M
K
G

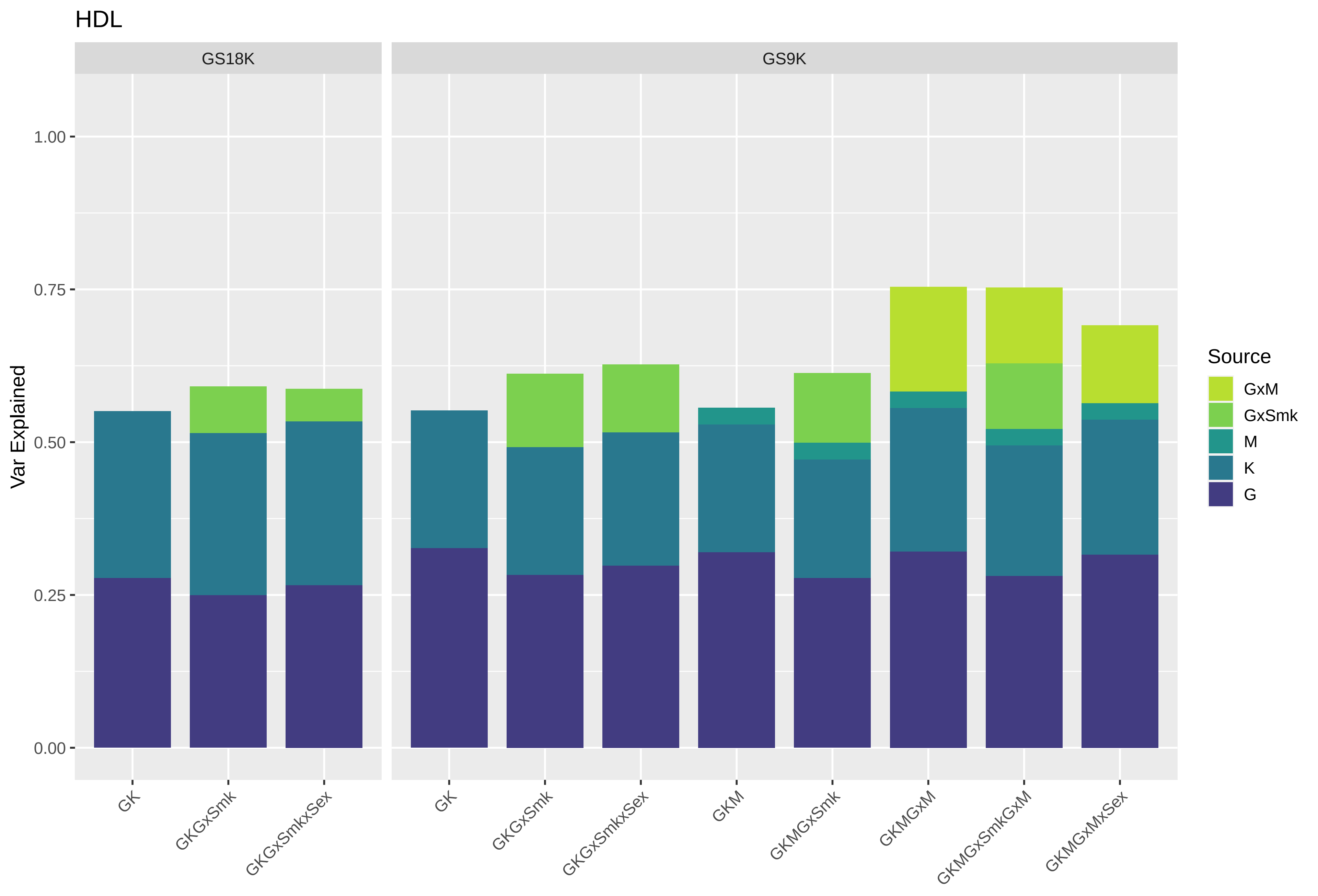

HDL
GS18K
GS9K
Var Explained
1.00
0.75
0.50
0.25
0.00
GK
GKGxSmk
GKGxSmkxSex
GK
GKGxSmk
GKGxSmkxSex
GKM
GKMGxSmk
GKMGxM
GKMGxSmkGxM
GKMGxMxSex
Source
GxM
GxSmk
M
K
G

**Supplementary Figure 2. Proportion of trait variation explained by Genome-by-Smoking interactions across all cohorts and sub-cohorts in each of the eight traits studied.** The plot shows the proportion of trait variance (the bars represent standard errors) explained by the genome-by-smoking interaction (x-axis) in the mixed model analyses across cohorts (y-axis). Panels from top to bottom represent cohorts: Generation Scotland (GS), UK Biobank (UKB), UK Biobank females (UKB_F) and UK Biobank males (UKB_M). Blue coloured data points show sub-cohort results (GS18K and UKB subgroups G1-G4), green coloured data points show meta-analyses of the corresponding panel sub-cohorts.

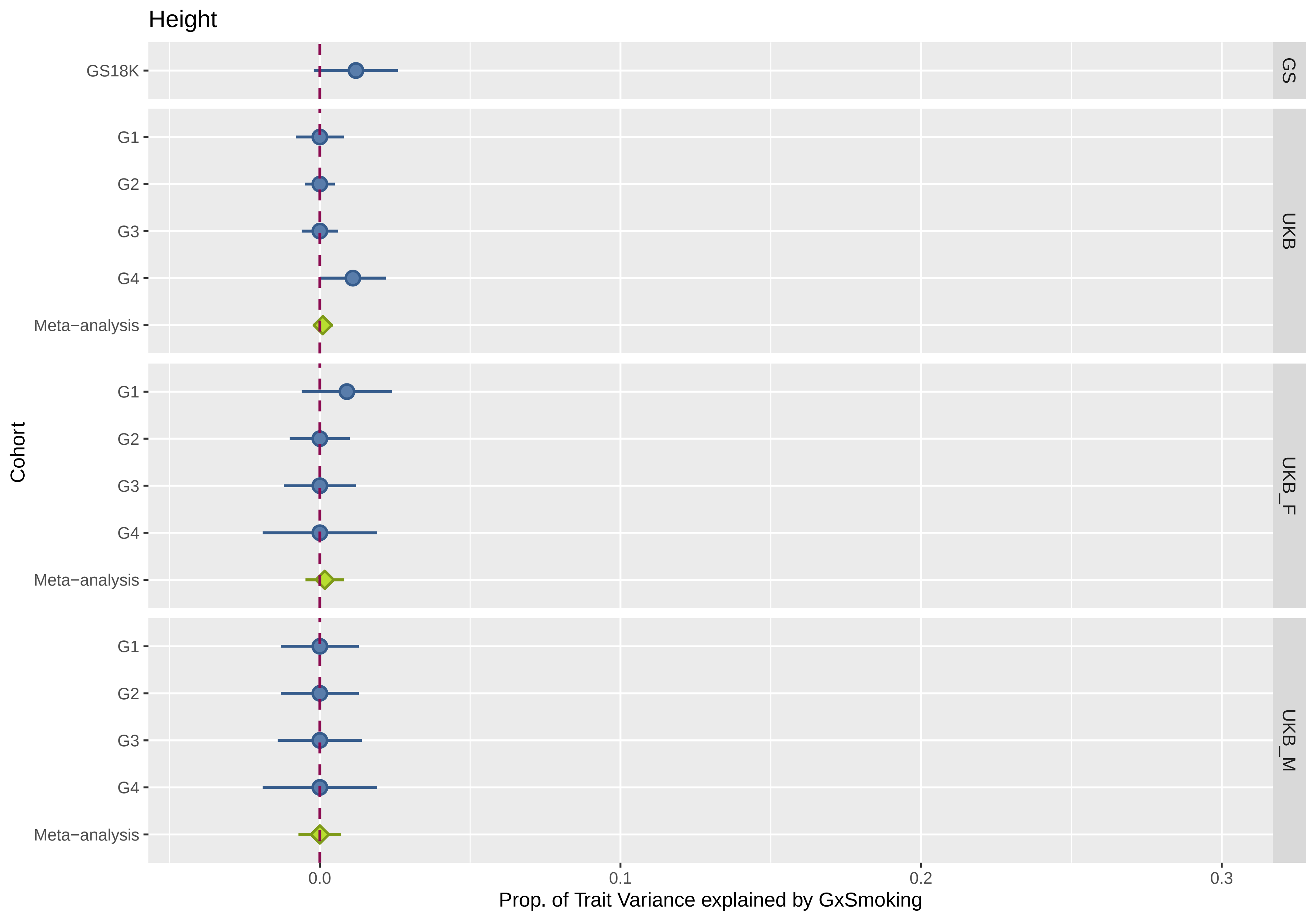

Height
GS18K
GS
G1
G2
G3
G4
Meta−analysis
UKB
G1
G2
G3
G4
Meta−analysis
UKB_F
G1
G2
G3
G4
Meta−analysis
UKB_M
Cohort
0.0
0.1
0.2
0.3
Prop. of Trait Variance explained by GxSmoking

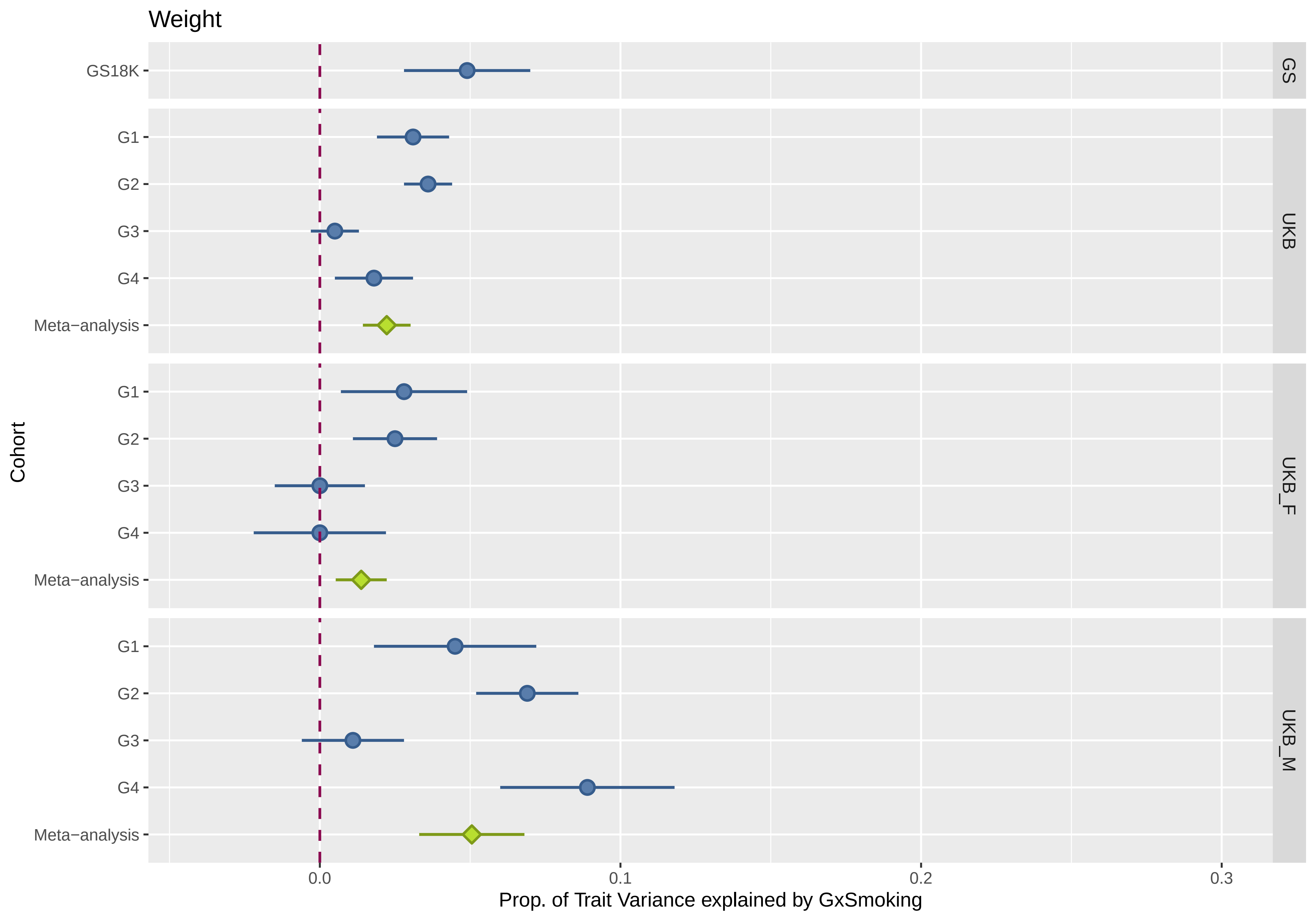
Weight
GS18K
GS
G1
G2
G3
G4
Meta−analysis
UKB
G1
G2
G3
G4
Meta−analysis
UKB_F
G1
G2
G3
G4
Meta−analysis
UKB_M
Cohort
0.0
0.1
0.2
0.3
Prop. of Trait Variance explained by GxSmoking

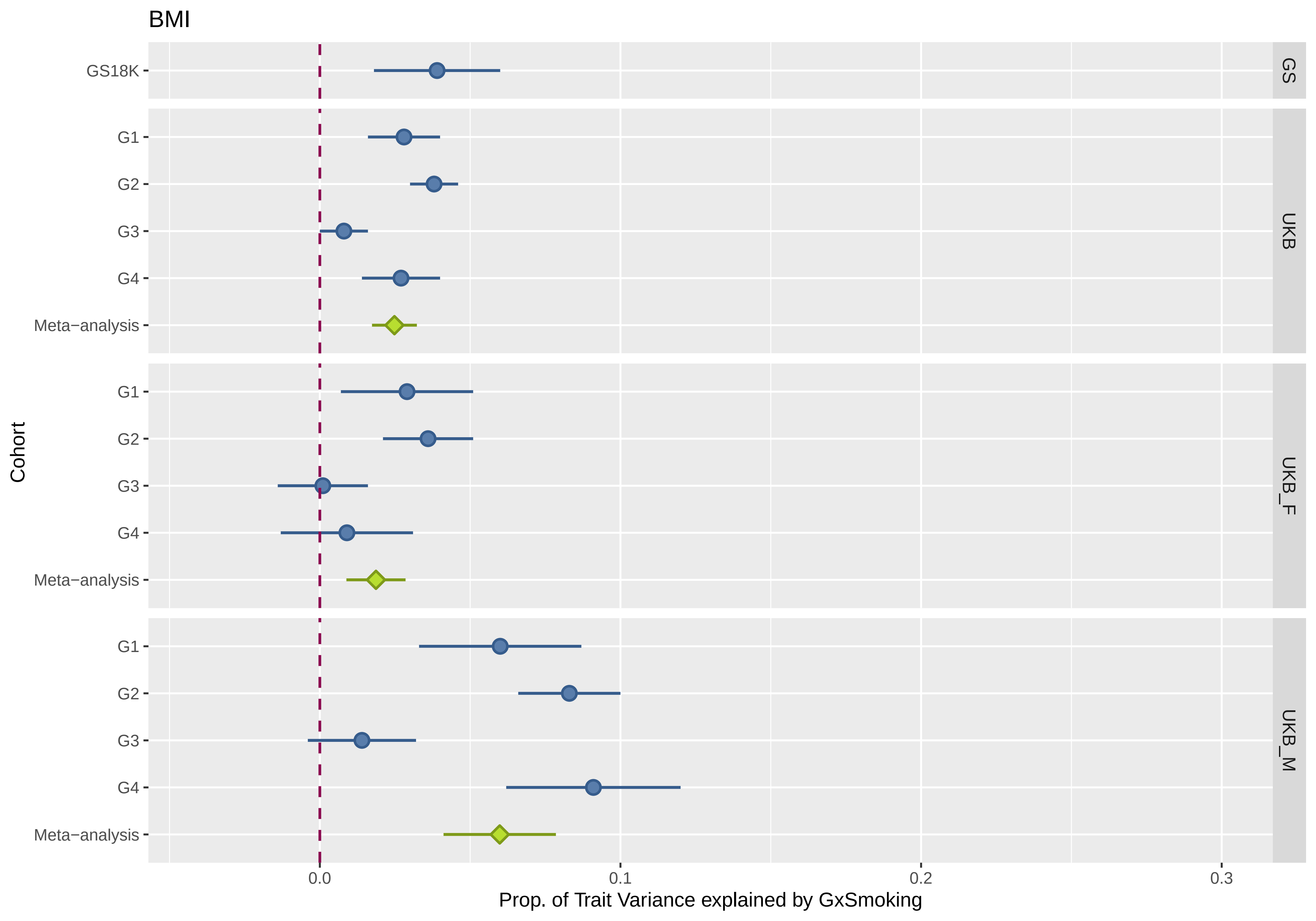

BMI
GS18K
G1
G2
G3
G4
Meta−analysis
G1
G2
G3
G4
Meta−analysis
G1
G2
G3
G4
Meta−analysis
GS
UKB
UKB_F
UKB_M
Cohort
0.0
0.1
0.2
0.3
Prop. of Trait Variance explained by GxSmoking

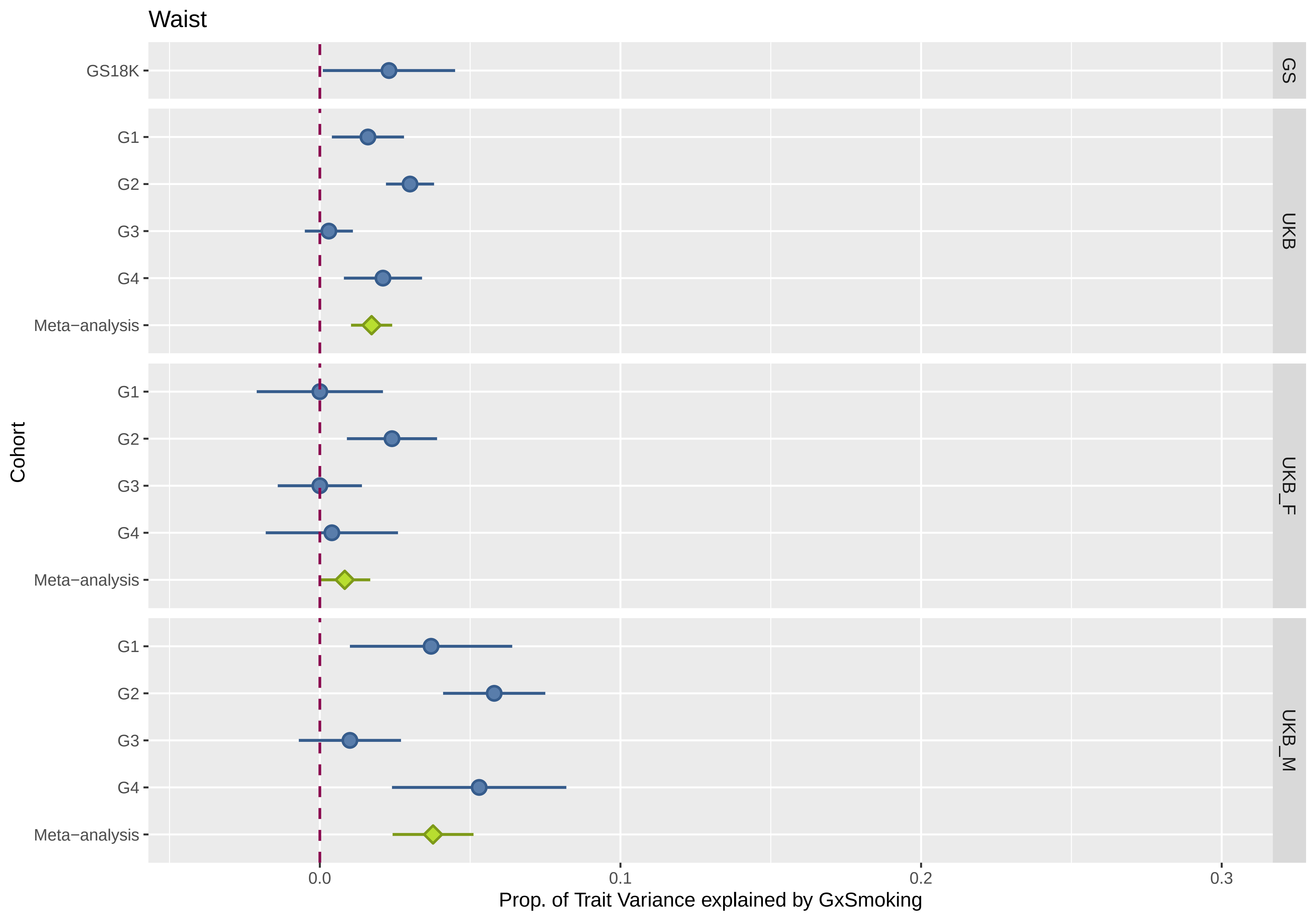

Waist
GS18K
G1
G2
G3
G4
Meta−analysis
G1
G2
G3
G4
Meta−analysis
G1
G2
G3
G4
Meta−analysis
GS
UKB
UKB_F
UKB_M
Cohort
0.0
0.1
0.2
0.3
Prop. of Trait Variance explained by GxSmoking

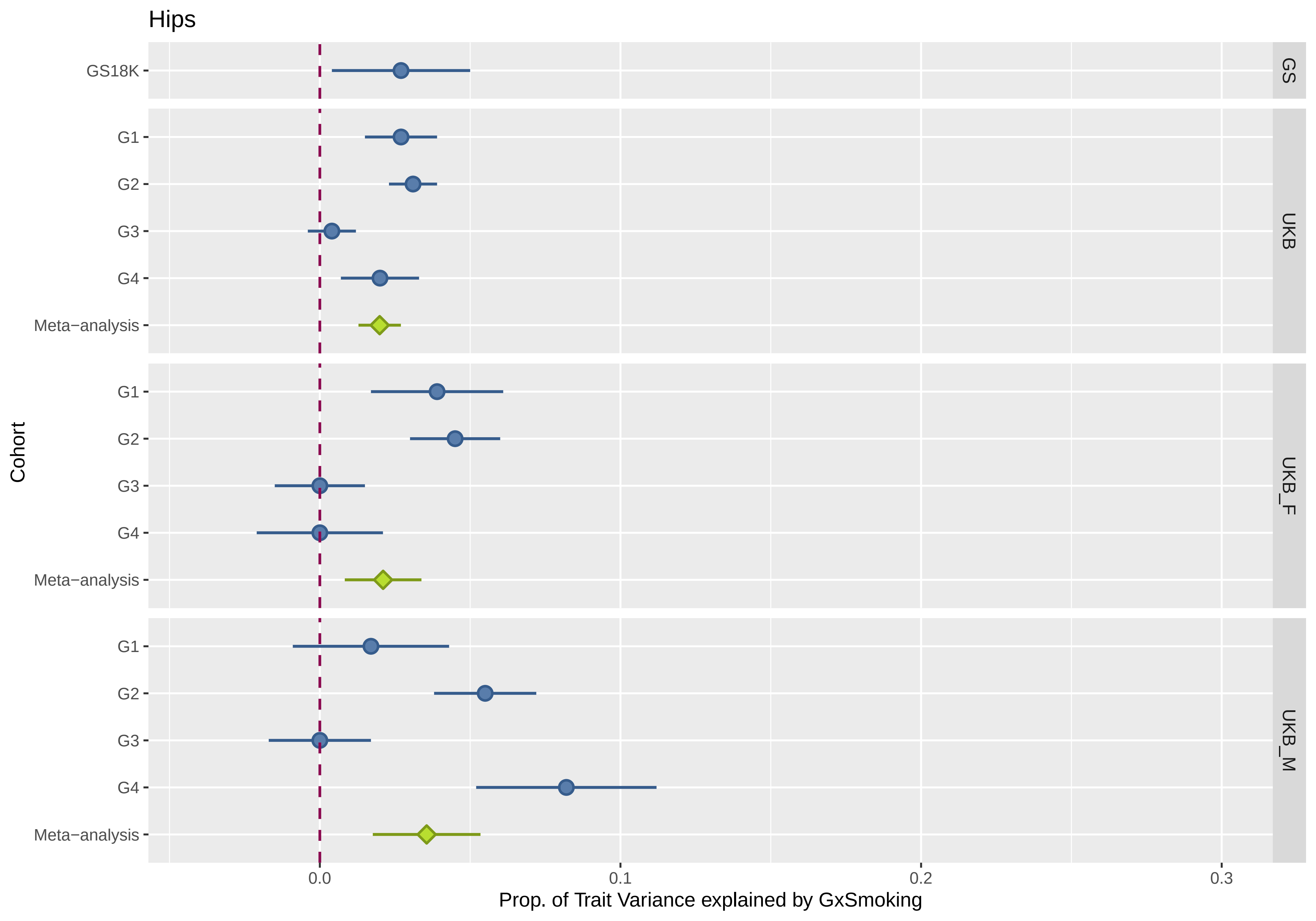

Hips
GS18K
G1
G2
G3
G4
Meta−analysis
G1
G2
G3
G4
Meta−analysis
G1
G2
G3
G4
Meta−analysis
GS
UKB
UKB_F
UKB_M
Cohort
0.0
0.1
0.2
0.3
Prop. of Trait Variance explained by GxSmoking

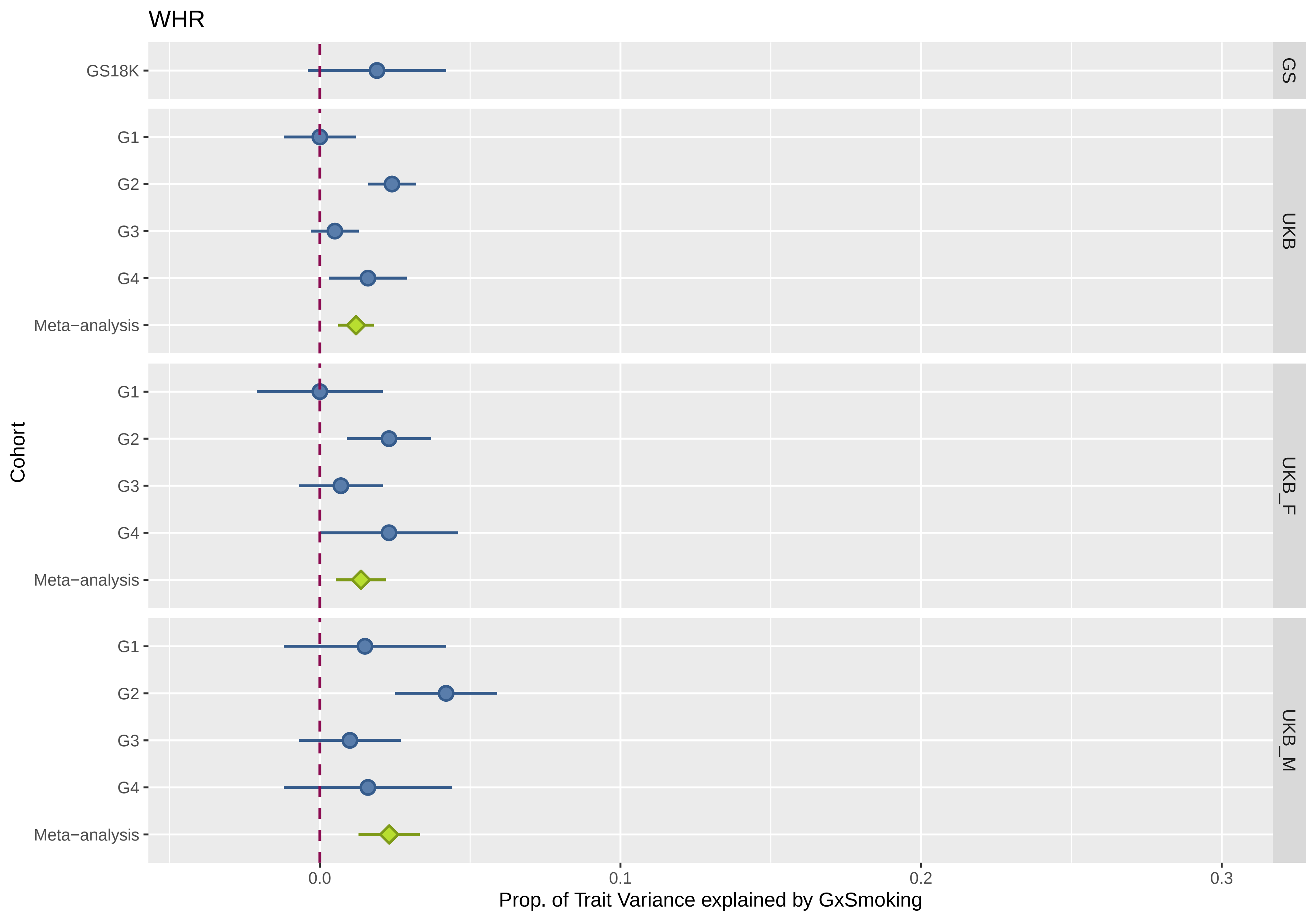

WHR
GS18K
G1
G2
G3
G4
Meta−analysis
G1
G2
G3
G4
Meta−analysis
G1
G2
G3
G4
Meta−analysis
GS
UKB
UKB_F
UKB_M
Cohort
0.0
0.1
0.2
0.3
Prop. of Trait Variance explained by GxSmoking

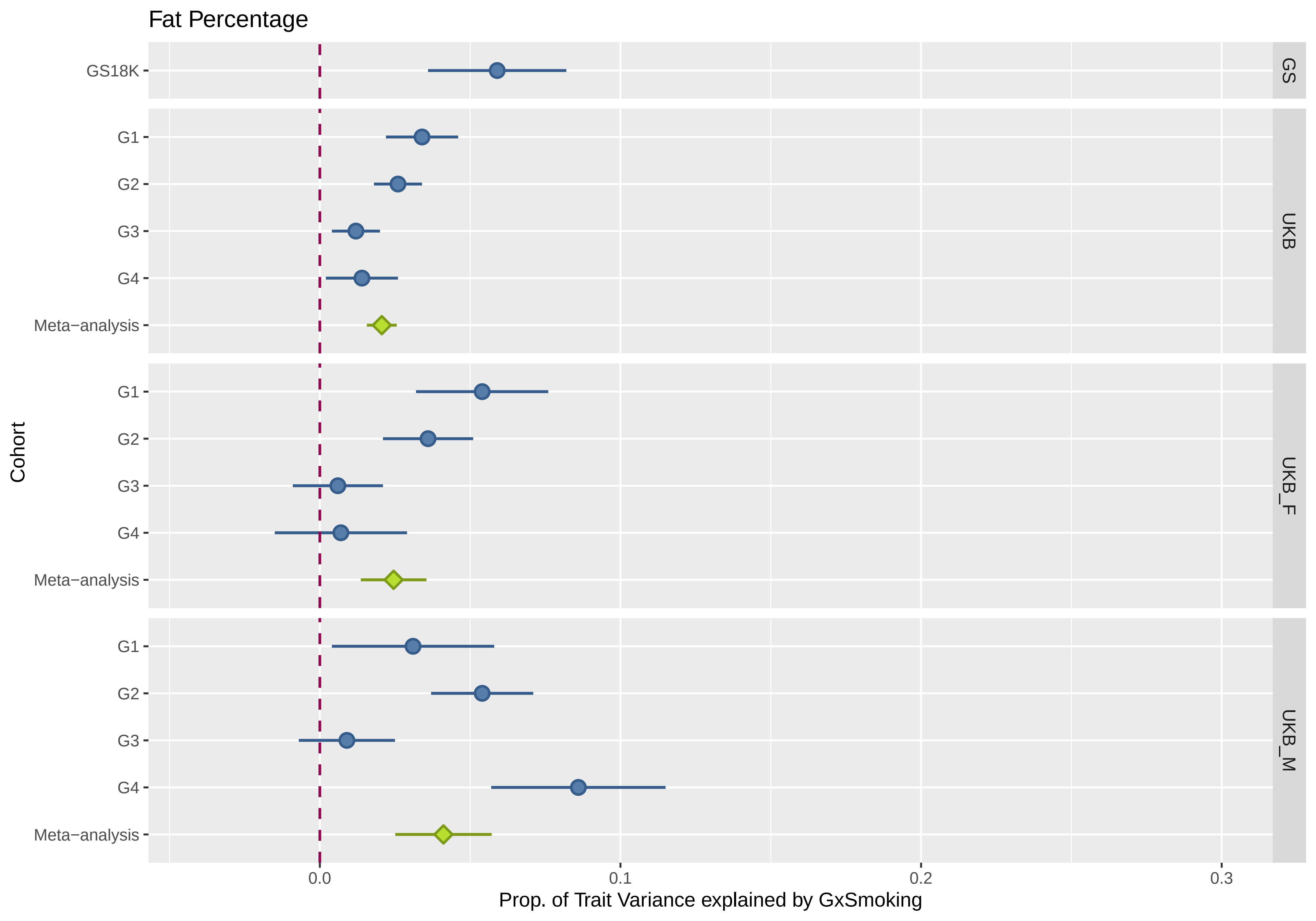

Fat Percentage
GS18K
G1
G2
G3
G4
Meta−analysis
G1
G2
G3
G4
Meta−analysis
G1
G2
G3
G4
Meta−analysis
GS
UKB
UKB_F
UKB_M
Cohort
0.0
0.1
0.2
0.3
Prop. of Trait Variance explained by GxSmoking

**Supplementary Figure 3. Proportion of trait variation explained by Genome-by-Smoking-by-Sex interactions across all cohorts and sub-cohorts in each of the eight traits studied.** The plot shows the proportion of BMI variance (the bars represent standard errors) explained by the genome-by-smoking-by-sex interaction (x-axis) in the mixed model analyses across cohorts (y-axis). Panels from top to bottom represent cohorts: Generation Scotland (GS), UK Biobank (UKB), UK Biobank females (UKB_F) and UK Biobank males (UKB_M). Blue coloured data points show sub-cohort results (GS18K and UKB subgroups G1-G4), green coloured data points show meta-analyses of the corresponding panel sub-cohorts.

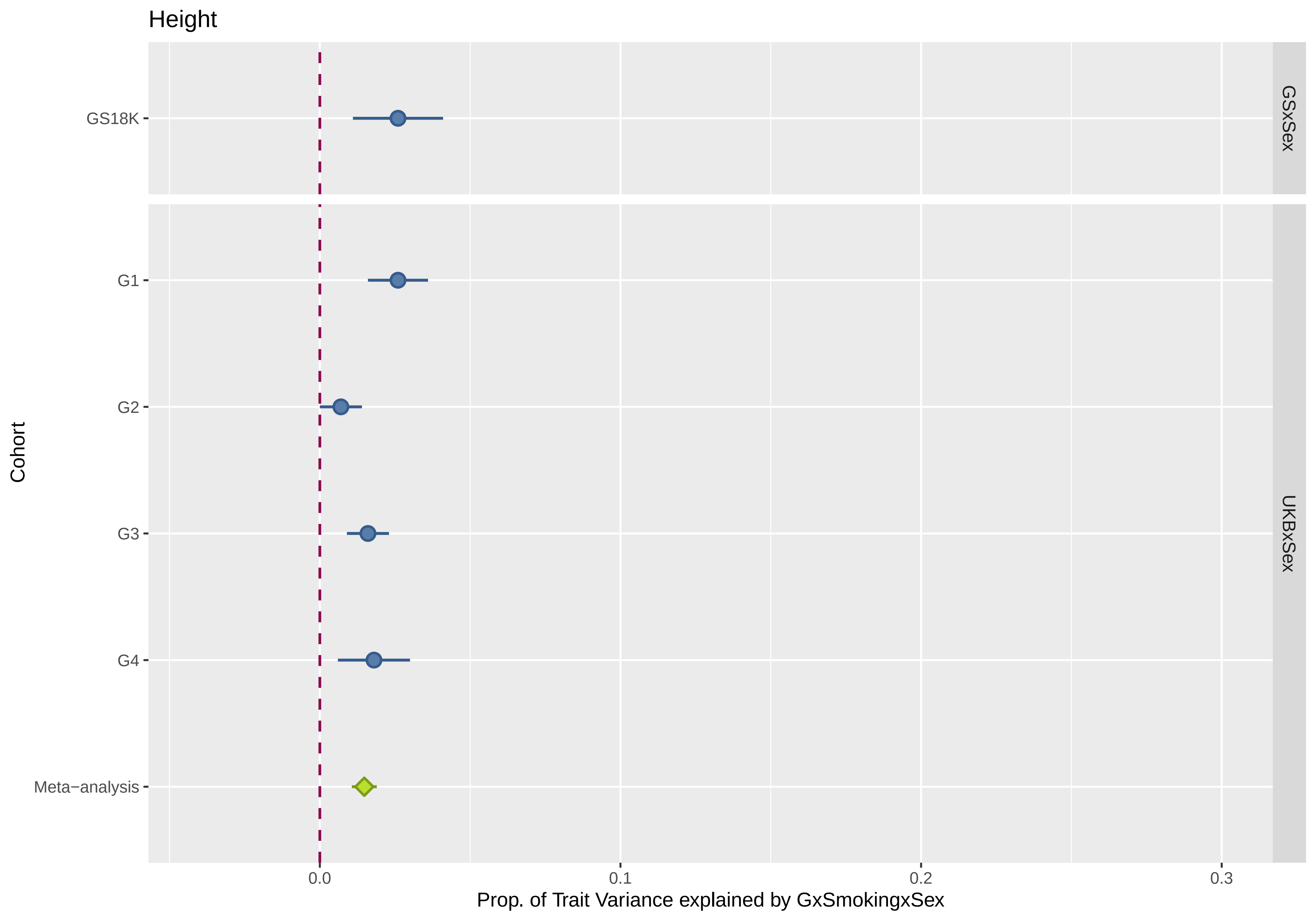
Height
GS18K
G1
G2
G3
G4
Meta−analysis
Cohort
GSxSex
UKBxSex
0.0
0.1
0.2
0.3
Prop. of Trait Variance explained by GxSmokingxSex

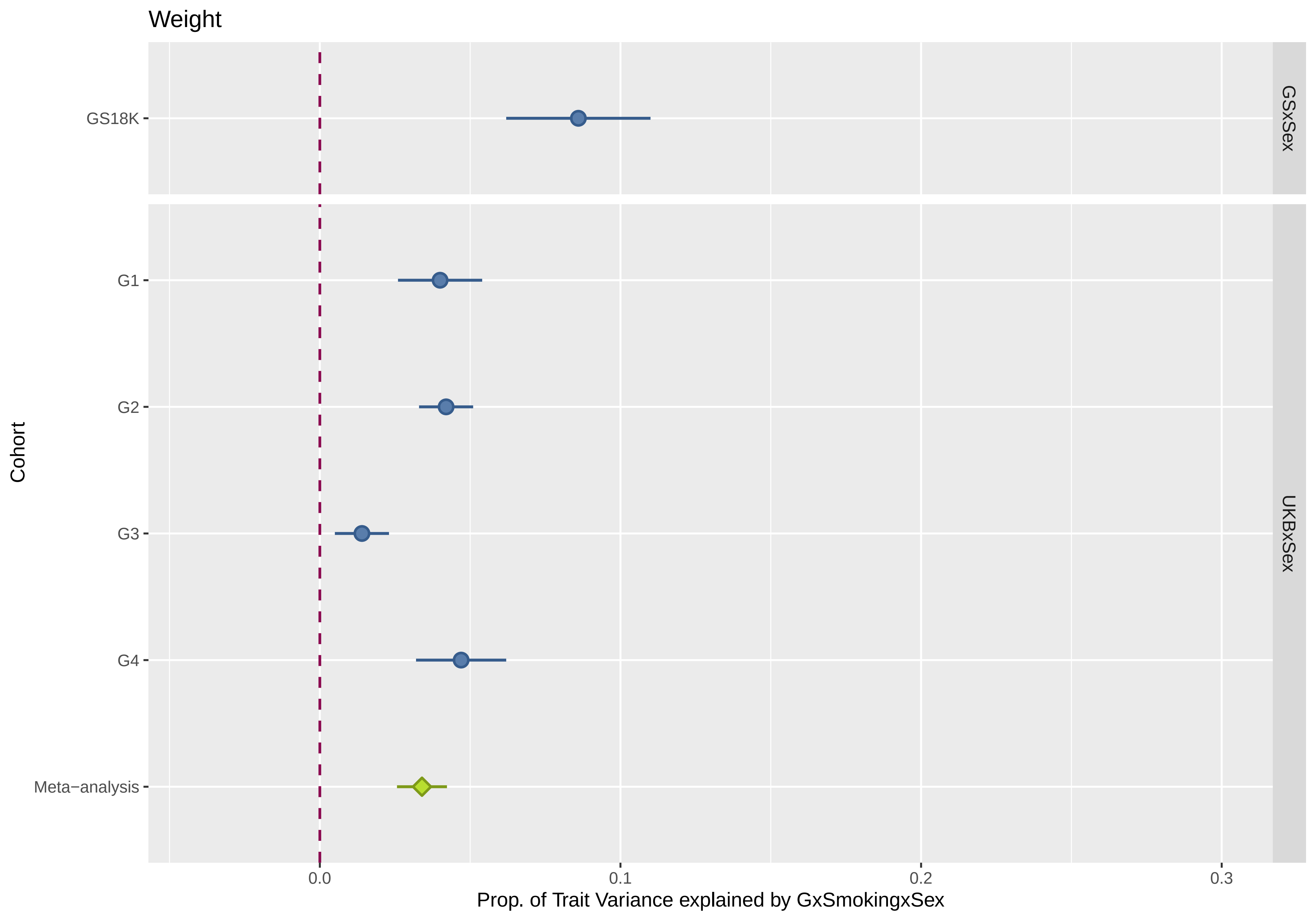
Weight
GS18K
G1
G2
G3
G4
Meta−analysis
Cohort
GSxSex
UKBxSex
0.0
0.1
0.2
0.3
Prop. of Trait Variance explained by GxSmokingxSex

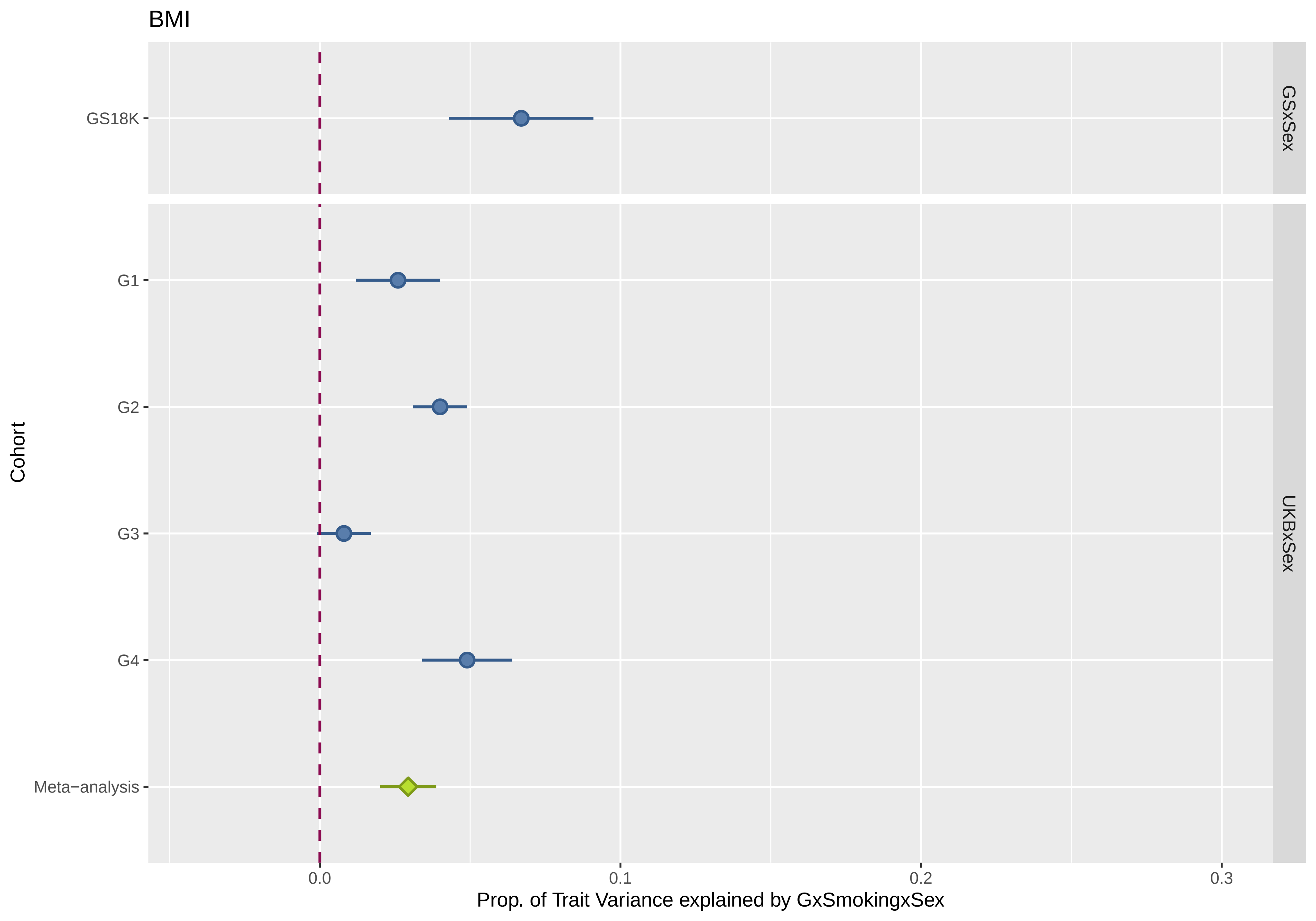

BMI
GS18K
G1
G2
G3
G4
Meta−analysis
Cohort
GSxSex
UKBxSex
0.0
0.1
0.2
0.3
Prop. of Trait Variance explained by GxSmokingxSex

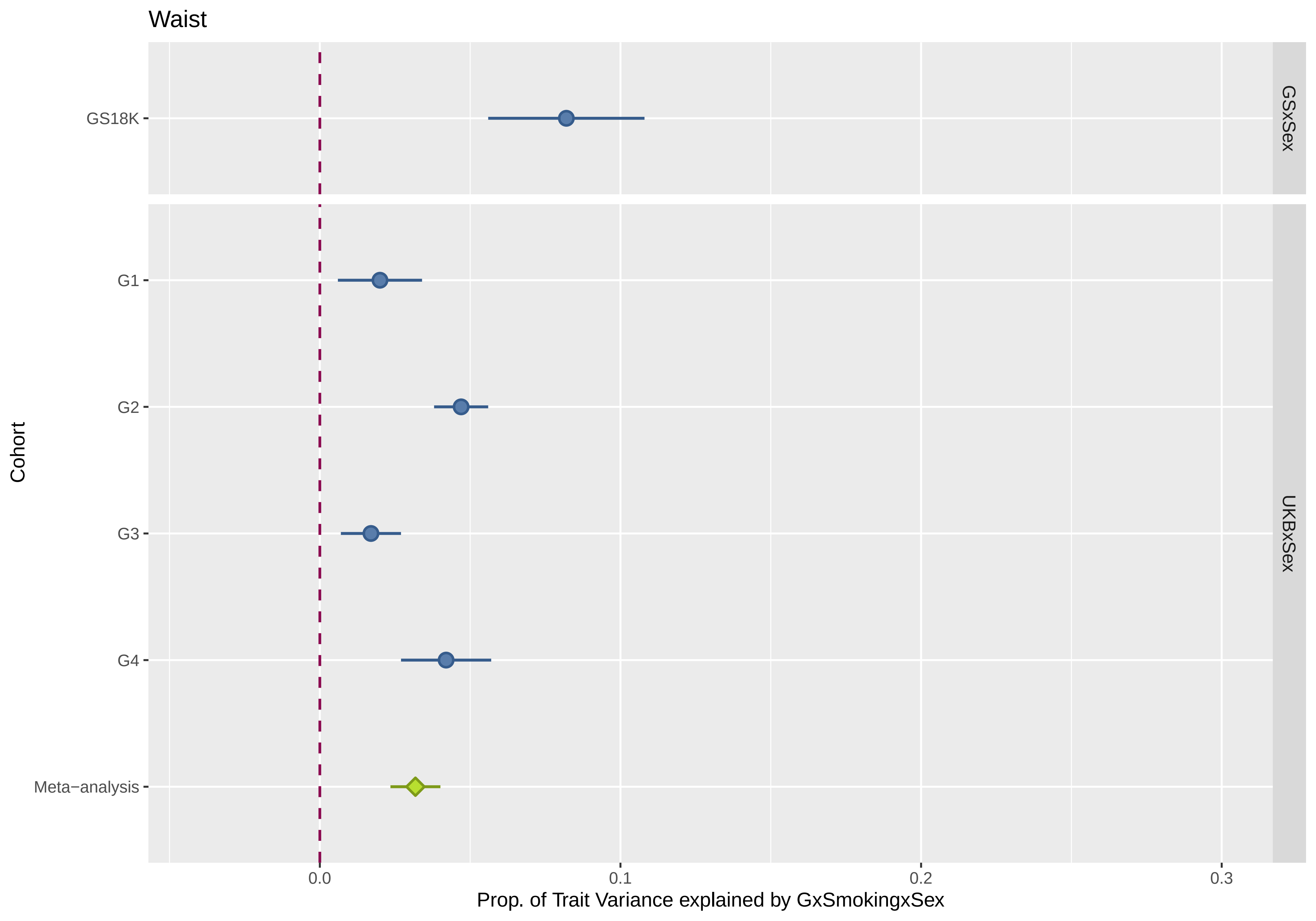

Waist
GS18K
G1
G2
G3
G4
Meta−analysis
Cohort
GSxSex
UKBxSex
0.0
0.1
0.2
0.3
Prop. of Trait Variance explained by GxSmokingxSex

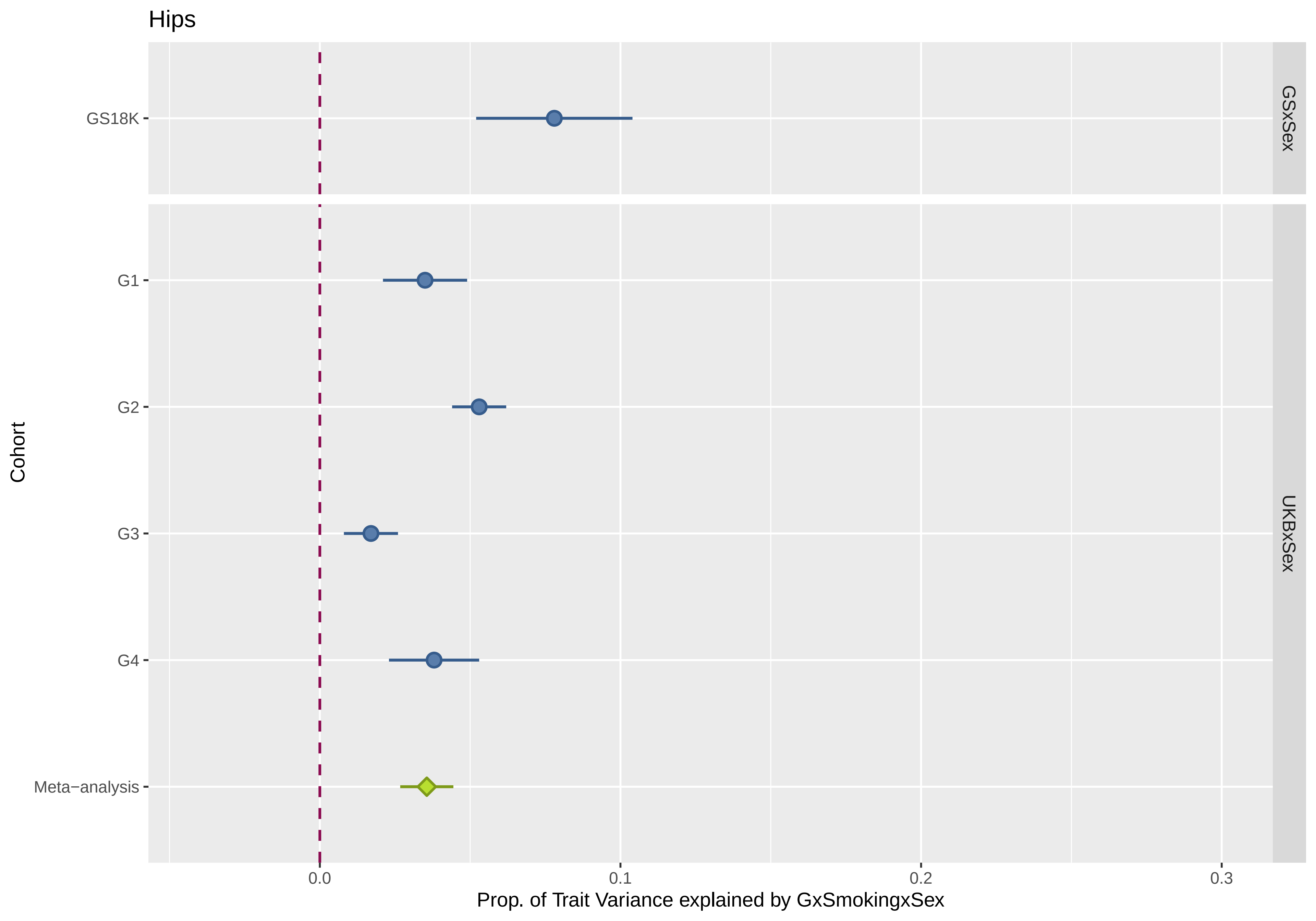

Hips
GS18K
G1
G2
G3
G4
Meta−analysis
Cohort
GSxSex
UKBxSex
0.0
0.1
0.2
0.3
Prop. of Trait Variance explained by GxSmokingxSex

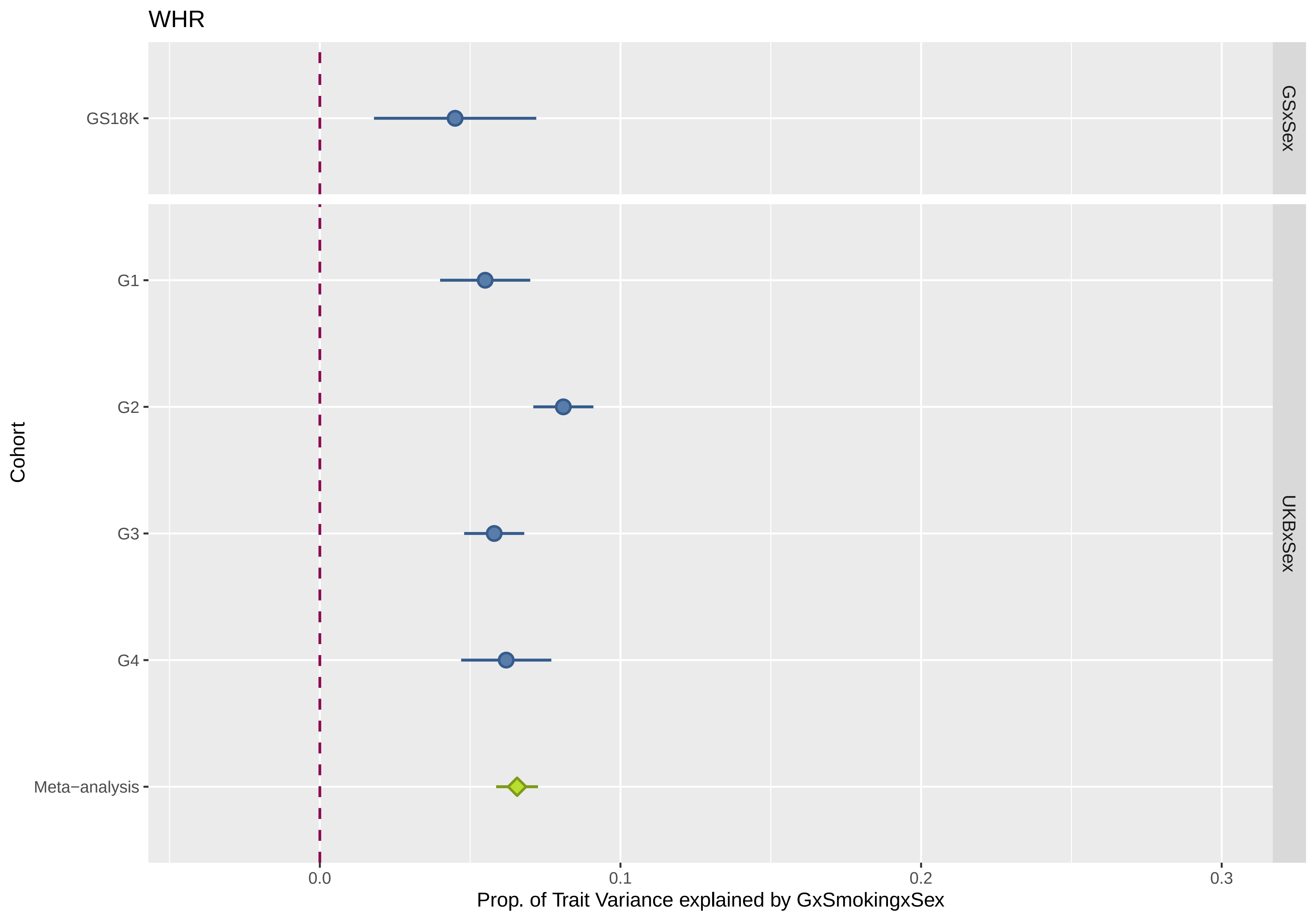
WHR
GS18K
G1
G2
G3
G4
Meta−analysis
Cohort
GSxSex
UKBxSex
0.0
0.1
0.2
0.3
Prop. of Trait Variance explained by GxSmokingxSex

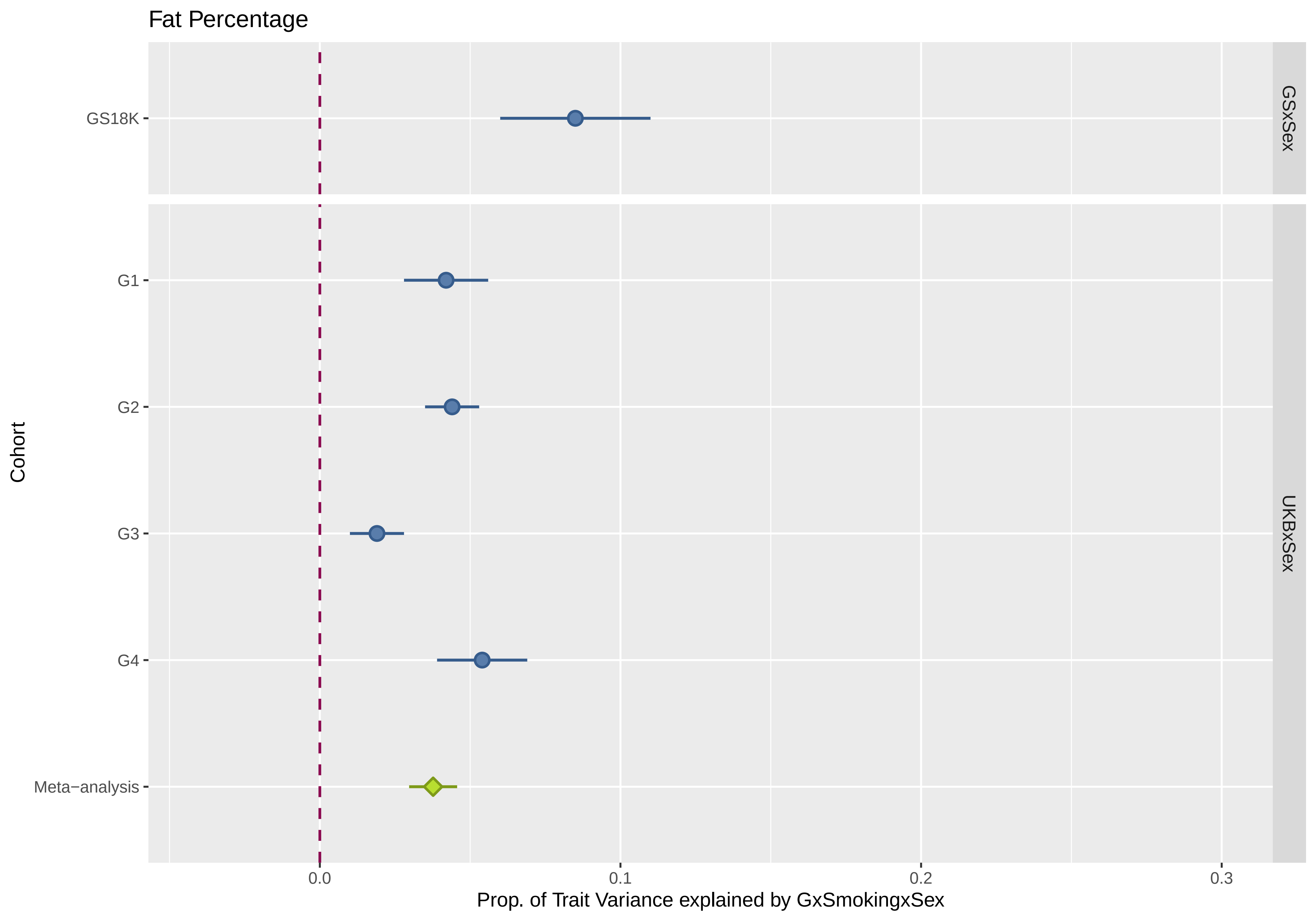

Fat Percentage
GS18K
G1
G2
G3
G4
Meta−analysis
Cohort
GSxSex
UKBxSex
0.0
0.1
0.2
0.3
Prop. of Trait Variance explained by GxSmokingxSex
