## Supplementary Text 1 for "Genome-wide methylation data improves dissection of the effect of smoking on body mass index"

### 1. Simulation of Phenotypes

In order to show that our models provide accurate estimates we performed a series of simulations. We simulated different sets of phenotypes for each of the 9,537 individuals in GS9K using their real genotypic and methylation profiles. These simulated phenotypes were generated by combining varying proportions of simulated effects associated with SNP variation, methylation variation and the interaction of the two.

#### Genetic effects

The genetic effects were simulated following Xia et al (2016). We used real information of genotyped markers located in even chromosomes to reduce computational burden ($N_{SNP}$ = 230,954). In each replicate, we randomly selected 1,000 causal SNPs to assign them effects from an exponential distribution with lambda parameter:

$$\lambda = \sqrt{\frac{4 * mean(pq) * 1000}{h_G^2}}$$

Where mean(*pq*) is the mean of the product of the minor allele frequencies multiplied by the major allele frequencies for these loci (0.194 here). $h^2_g$ is the proportion of the variance of the simulated phenotype that is driven by genetic effects.

The vector of weighted summed effects for those loci across individuals (*g*) was calculated as:

$$g = \sum_{i=1}^{1000} a_i x_i$$

where $a_i$ is the effect size of the reference allele in SNP *i* and $x_i$ is the vector of allelic dosages for SNP *i*. We transformed *g* to a normal distribution with mean equal to 0 and variance equal to $h^2_g$.

#### Methylation effects

The 62 smoking-associated CpG sites (Supplementary Table 9) contributed to the methylation effects. We simulated their effect sizes from a normal distribution $N(0, h_M^2)$. $h^2_M$ is the proportion of the variance of the simulated phenotype that is driven by methylation effects.

The vector of summed effect for those loci (*m*) was calculated as:

$$m = \sum_{j=1}^{62} b_j w_j$$

where $b_i$ is the effect size of CpG site *j* and $w_j$ is the vector of methylation levels for CpG site *j*. We transformed *m* to a normal distribution with mean equal to 0 and variance equal to $h^2_M$.

#### Genome-by-Methylation effects

The 62 smoking-associated CpG sites (Supplementary Table 9) contributed to the genome-by-methylation effects. For each CpG site we selected a random set of 1000 SNPs and assigned them effect sizes from a normal distribution $N(0, h_{GxM}^2)$. $h^2_{GxM}$ is the proportion of the variance of the simulated phenotype that is driven by genome-by-methylation interaction effects.

The vector of weighted summed effects for those loci (*gxm*) was calculated as:

$$gxm = \sum_{j=1}^{62}\left(\sum_{i=1}^{1000} c_i * x_i\right) * w_j$$

where $c_i$ is the effect size of the interaction with SNP *i*, $x_i$ is the vector of allelic dosages for SNP *i*, and $w_j$ is the vector of methylation levels for CpG site *j*. We transformed *gxm* to a normal distribution with mean equal to 0 and variance equal to $h^2_{GxM}$.

Both methylation and genome-by-methylation effects are derived from a small subset of 62 CpGs. Since the interaction effects are computed as the product of the methylation level at those 62 CpGs by a group of genetic effects part of these effects could be considered mean effects and will be correlated with the methylation effects. This could be reflected in the estimation of variance parameters (i.e., $h^2_{gxm}$ is actually smaller than the simulated value).

### Residual effects and construction of the phenotypes

In each scenario, each component ($h^2_g$, $h^2_m$, $h^2_{gxm}$) contributed a proportion of the variance explained (0-50%). A residual effect ($\varepsilon$) was simulated with values obtained from a normal distribution $N(0, e^2)$, where $e^2$ represents the proportion of variance unexplained by the three sources combined in each of the scenarios:

$$e^2 = 1 - h^2_G - h^2_M - h^2_{GxM}$$

The final phenotypes were constructed by adding the previously described components and had an expected mean and variance of 0 and 1, respectively.

$$pheno = g + m + gxm + \varepsilon$$

### Scenarios

The four scenarios simulated are shown in Table ST1.1. For each scenario we performed a total of 50 replicates. The scenarios aimed to mimic the results observed in the real data by simulating phenotypic contributions similar to those estimated for BMI (Supplementary Table 8).

| COMPONENT | SCENARIO 1 | SCENARIO 2 | SCENARIO 3 | SCENARIO 4 |
|---|---|---|---|---|
| G | 0.50 | 0.50 | 0.50 | 0.00 |
| M | 0.00 | 0.05 | 0.05 | 0.00 |
| GxM | 0.00 | 0.00 | 0.25 | 0.25 |
| Res | 0.50 | 0.45 | 0.20 | 0.75 |

**Table ST1.1.** Scenarios tested in the simulations. The table shows the proportion of the phenotypic variance contributed by each component in each of the four simulated scenarios.

### 2. Model testing in the Simulated Phenotypes

For each scenario and replicate, we fitted a series of mixed models with different combinations of similarity matrices to estimate the variance explained by each component as in the real data. For consistency with the phenotype simulations, the genomic similarity matrix was created using the same subset of 230,954 SNPs used and similarly, a corresponding genome-by-methylation matrix was derived from this 230K genomic matrix. The methylation similarity matrix was the same one used with the real data, derived from the 62 smoking associated CpG sites. (For more information see main methods section).

### 3. Simulation Results

#### Scenario 1.

In the first scenario, we examined the performance of our analysis models when the simulated phenotypes consisted only of a genetic effect (and residual variation). Summary results of the 50 replicates of this scenario are shown in Figure ST1.1.

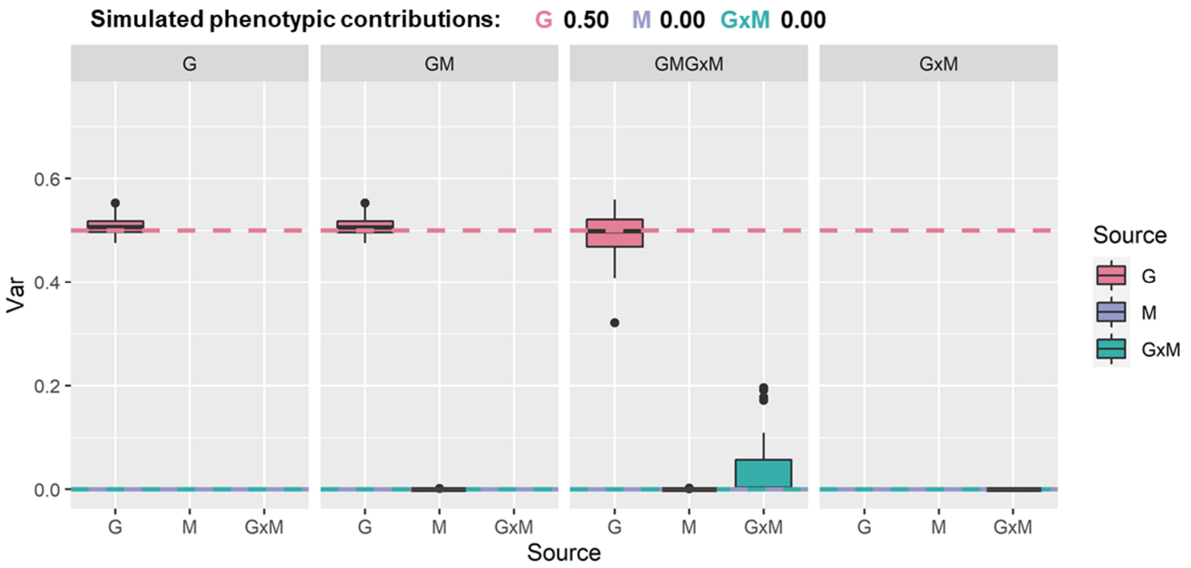


**Figure ST1.1. Results of the variance component analyses for simulated scenario 1.** Estimates of the proportion of the phenotypic variance (y-axis) explained by each effect fitted in the analytical model (x-axis and colours). Boxes represent results from 50 replicates (medians and inter-quantile ranges). A black bar at 0.0 indicates that all replicates produced an estimate of zero for the relevant component. Panels show results for models fitted including different combinations of the variance components as defined in the panel headings. Dashed lines represent the simulated proportion of the phenotypic variance contributed by each component indicated by colours in the legend. G: Genomic, M: Smoking associated methylation, GxM: Genome-by-Methylation.

The results show that the genomic matrix (G) provides unbiased estimates of the simulated variation across all models. The smoking-associated methylation matrix (M) does not capture any variation, as expected. The genome-by-methylation matrix (GxM) provides some positive estimates, but these were only significantly different from zero in 2 out of 50 replicates (consistent with a significance threshold of 5%). When testing a model without G and M components, GxM does not capture any variation (estimates from all replicates were zero).

Scenario 2.

In the second scenario, we examined the performance of our models when a genetic and a methylation effect (and residual variation) contributed to the simulated phenotype. Summary results of the 50 replicates of this scenario are shown in Figure ST1.2.

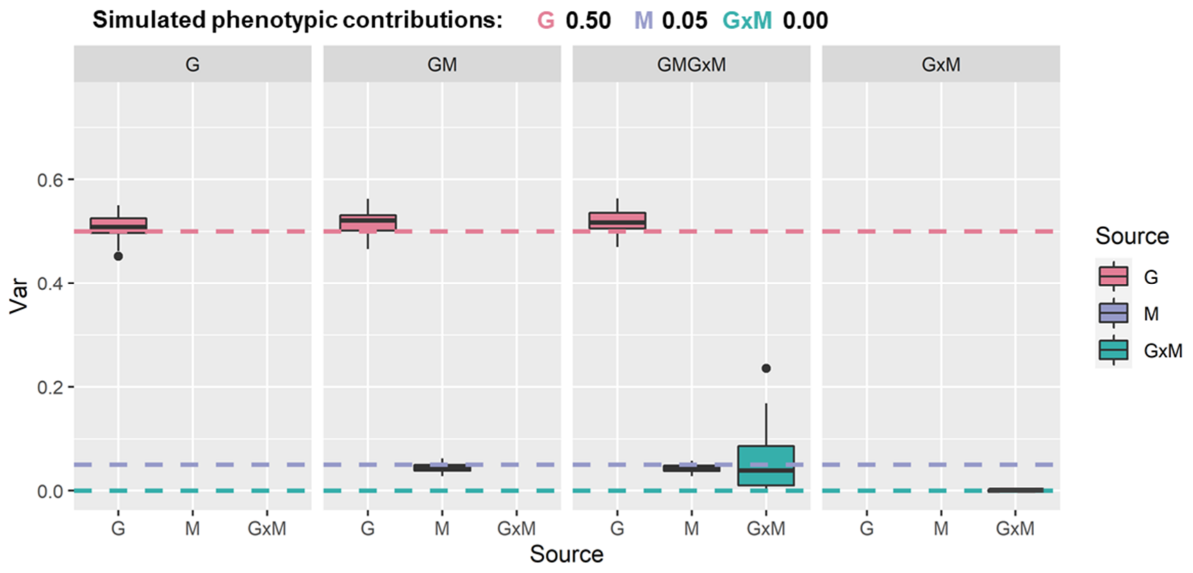


**Figure ST1.2. Results of the variance component analyses for simulated scenario 2.** Estimates of the proportion of the phenotypic variance (y-axis) explained by each effect fitted in the analytical model (x-axis and colours). Boxes represent results from 50 replicates (medians and inter-quantile ranges). A black bar at 0.0 indicates that all replicates produced an estimate of zero for the relevant component. Panels show results for models fitted including different combinations of the variance components as defined in the panel headings. Dashed lines represent the simulated proportion of the phenotypic variance contributed by each component indicated by colours in the legend. G: Genomic, M: Smoking associated methylation, GxM: Genome-by-Methylation.

The results show that in this scenario, both G and M provide unbiased estimates of the simulated variation across all models. As before, GxM provides some positive estimates (1 of 50 were significantly different from zero) and, when testing a model without G and M components, GxM cannot capture any significant variation.

Scenario 3.

In the third scenario, we examined the performance of our models when all components: genetic, methylation and genome-by-methylation effects (and residual variation) contributed to the simulated phenotype. Summary results of the 50 replicates of this scenario are shown in Figure ST1.3.

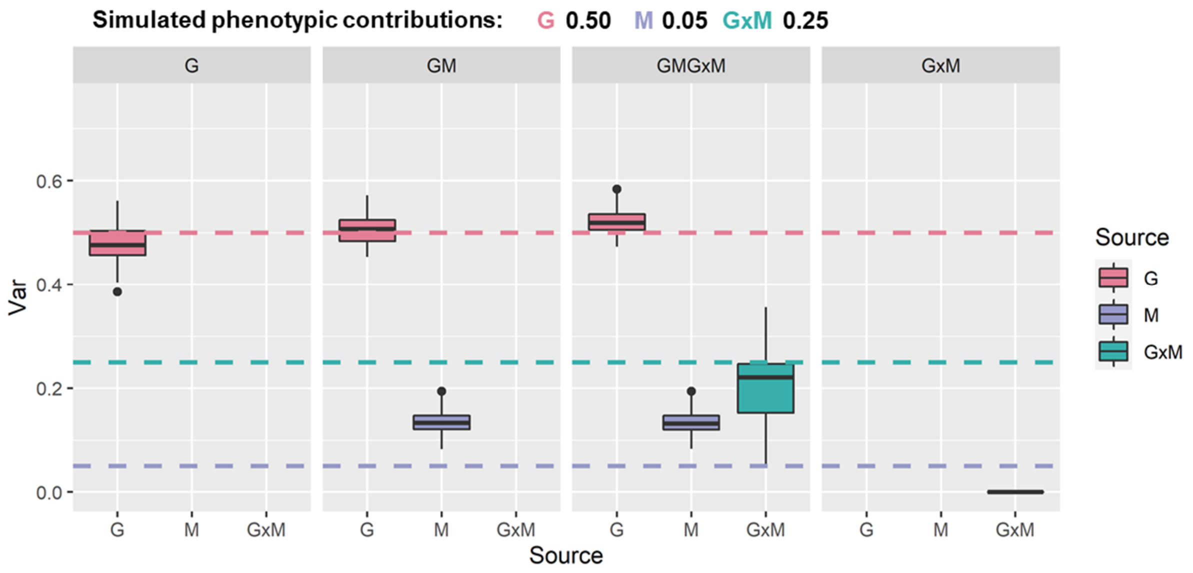


**Figure ST1.3. Results of the variance component analyses for simulated scenario 3**. Estimates of the proportion of the phenotypic variance (y-axis) explained by each effect fitted in the analytical model (x-axis and colours). Boxes represent results from 50 replicates (medians and inter-quantile ranges). A black bar at 0.0 indicates that all replicates produced an estimate of zero for the relevant component. Panels show results for models fitted including different combinations of the variance components as defined in the panel headings. Dashed lines represent the simulated proportion of the phenotypic variance contributed by each component indicated by colours in the legend. G: Genomic, M: Smoking associated methylation, GxM: Genome-by-Methylation.

The results show that in this scenario, G provides unbiased estimates of its simulated variation across all models. The proportion of the variance explained by M is larger than the effects attributed to it in the simulation, while the GxM captures less variation that was originally simulated. This may be driven by the way the interaction effects were created in the simulations. The effects of CpG sites were weighted by the individual's genotypes, but some of these effects are attributable to mean differences between CpG methylation levels, and therefore can be picked up by the M matrix. This is a limitation of this simulation since in this particular scenario, the proportions of variance attributable to M and GxM are not exactly 0.05 and 0.25. The average variance captured by the sum of M and GxM was 0.32, very closed to the 0.30 simulated (0.05 + 0.25). The variance captured by GxM was significantly different from zero in 37/50 replicates when it is modelled jointly with the other components. When modelled without any other component, GxM was not able to capture any variance suggesting that without adjusting to the other effects GxM variation is difficult to estimate (consistent with the way the GxM was derived in the simulations).

Scenario 4.

In the final scenario, we examined the performance of our models when only genome-by-methylation effects (and residual variation) contributed to the simulated phenotype. Summary results of the 50 replicates of this scenario are shown in Figure ST1.4.

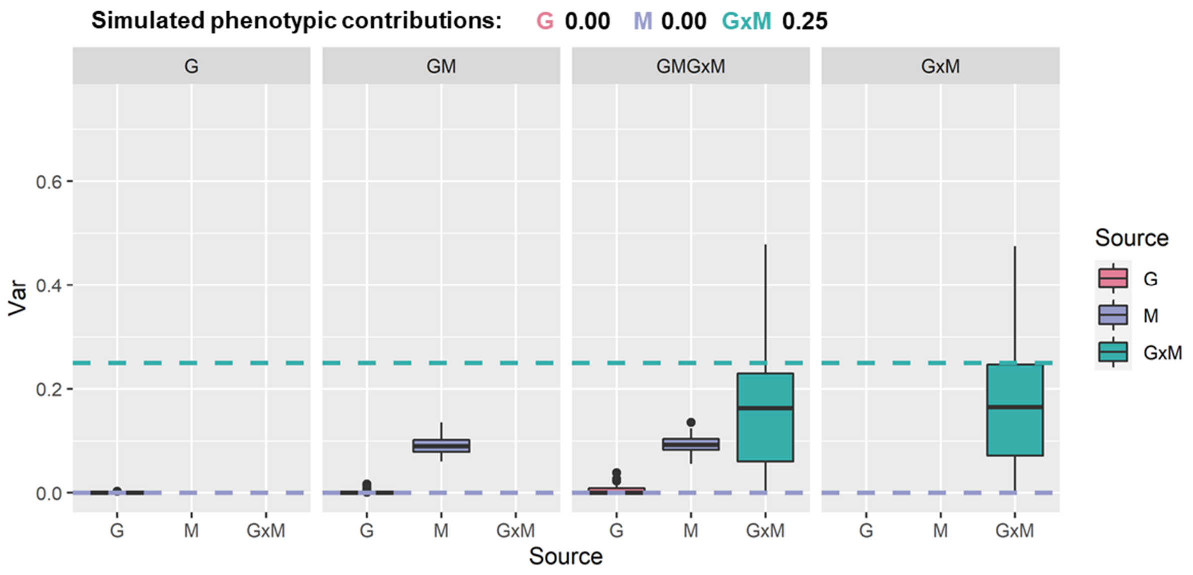


**Figure ST1.4. Results of the variance component analyses for simulated scenario 4.** Estimates of the proportion of the phenotypic variance (y-axis) explained by each effect fitted in the analytical model (x-axis and colours). Boxes represent results from 50 replicates (medians and inter-quantile ranges). A black bar at 0.0 indicates that all replicates produced an estimate of zero for the relevant component. Panels show results for models fitted including different combinations of the variance components as defined in the panel headings. Dashed lines represent the simulated proportion of the phenotypic variance contributed by each component indicated by colours in the legend. G: Genomic, M: Smoking associated methylation, GxM: Genome-by-Methylation.

As we observed in the previous scenario, these results confirm the relationship between the M and GxM effects. Some of the simulated GxM effects are captured by the M matrix confirming that part of these interaction effects are attributable to mean differences between individual’s methylation.
